# SHORT DISORDERED PEPTIDES ARE SUFFICIENT TO CONVERT PROTEINS INTO MECHANOSENSORS THAT RESPOND TO PHYSIOLOGICAL CELLULAR FORCES

**DOI:** 10.1101/2025.11.11.685779

**Authors:** Antuca Callejas-Marín, Walter Gonzalez, Tino Pleiner, Ting-Hao Huang, Aubrie de la Cruz, Ashutosh Pandey, Luis Sánchez-Guardado, Hamed Jafar-Nejad, Rebecca Voorhees, Carlos Lois

## Abstract

Mechanical forces regulate many biological processes and a handful of mechanosensor domains have been identified in proteins that respond to cellular forces. The Notch signaling pathway, a key regulator of cell communication processes ranging from animal development to cancer, is activated by mechanical forces generated by cell-cell interactions. The mechanosensing element of Notch is thought to be a 300 amino acid extracellular domain called negative regulatory region (NRR). Ligand binding exerts mechanical forces that are believed to partially unfold NRR, rendering it susceptible to cleavage by extracellular metalloproteases that leads to activation of signaling. However, some engineered notch-like receptors lacking the NRR can be activated upon ligand binding, although the mechanisms underlying this activation are not known. Here we observe that Notch molecules without the NRR, but with a 12 amino acid sequence present in the extracellular juxta-transmembrane domain (eJTMD) are also cleaved and activated upon interaction with their ligands in cultured cells and in transgenic animals. Furthermore, the ∼12 aa eJTMD from notch genes from other species (chicken, xenopus, zebrafish and *Drosophila*), from other unrelated transmembrane proteins (erbB4, N-CAM, and E-Cadherin), or multiple artificial sequences of ∼12 amino acids without any apparent structure or specific sequence can also convert proteins into ligand-dependent mechanosensors. These results indicate that short, disordered amino acid sequences that are commonly found in many proteins are capable of imparting mechanosensing capabilities into proteins, suggesting that mechanical force may regulate many more cellular processes than previously suspected.

## Introduction

Mechanical forces generated by cells regulate many physiological processes, ranging from selection of optimal antibodies by lymphocytes (Natkanski *et al*, 2013), sorting of chromosomes within the cell (Pavin & Tolić, 2021), and blood clotting in response to turbulence (Zhang *et al*, 2009b). Accordingly, cells have evolved specialized proteins capable of sensing these forces (Yusko & Asbury, 2014). One of the best studied cellular mechanosensors is notch, a protein involved in multiple developmental processes, whose dysregulation causes several diseases, most notably leukemias (McIntyre *et al*, 2020). Notch is a type I transmembrane protein that contains several domains with distinct functions (Kovall *et al*, 2017). Its extracellular domain (ECD) contains the ligand binding domain (LBD) and the negative regulatory region (NRR) (Kovall *et al*, 2017). The LBD is necessary to bind to notch’s ligands, delta, serrate and jagged. The NRR is a 300 aa domain that contains within it a site called S2 that can be cleaved by ubiquitous membrane-bound metalloproteases of the ADAM family (Lovendahl *et al*, 2018) (**Fig. 1A and B**). In the absence of ligand, the NRR is folded such that the S2 site is buried, and the ADAM metalloproteases cannot access it. It is thought that upon binding of notch to its ligands, the mechanical force produced by endocytosis of the ligand triggers local unfolding of the NRR and exposure of the S2 site, which is then cleaved by ADAM metalloproteases (Tousseyn *et al*, 2009). After S2 cleavage, the fragment of notch ECD upstream of S2 dissociates from the fragment downstream of it, such that only a short stump 12 aa long of notch ECD, its extracellular juxtatransmembrane domain (ejTMD), remains above the membrane. This shortening of the notch ECD triggers an intramembrane cleavage by the gamma-secretase cleavage in the so-called S3 site, freeing the notch ICD such that it reaches the nucleus and activates transcription (De Strooper et al., 1999) (**Fig. 1B**).

**Figure 1:**
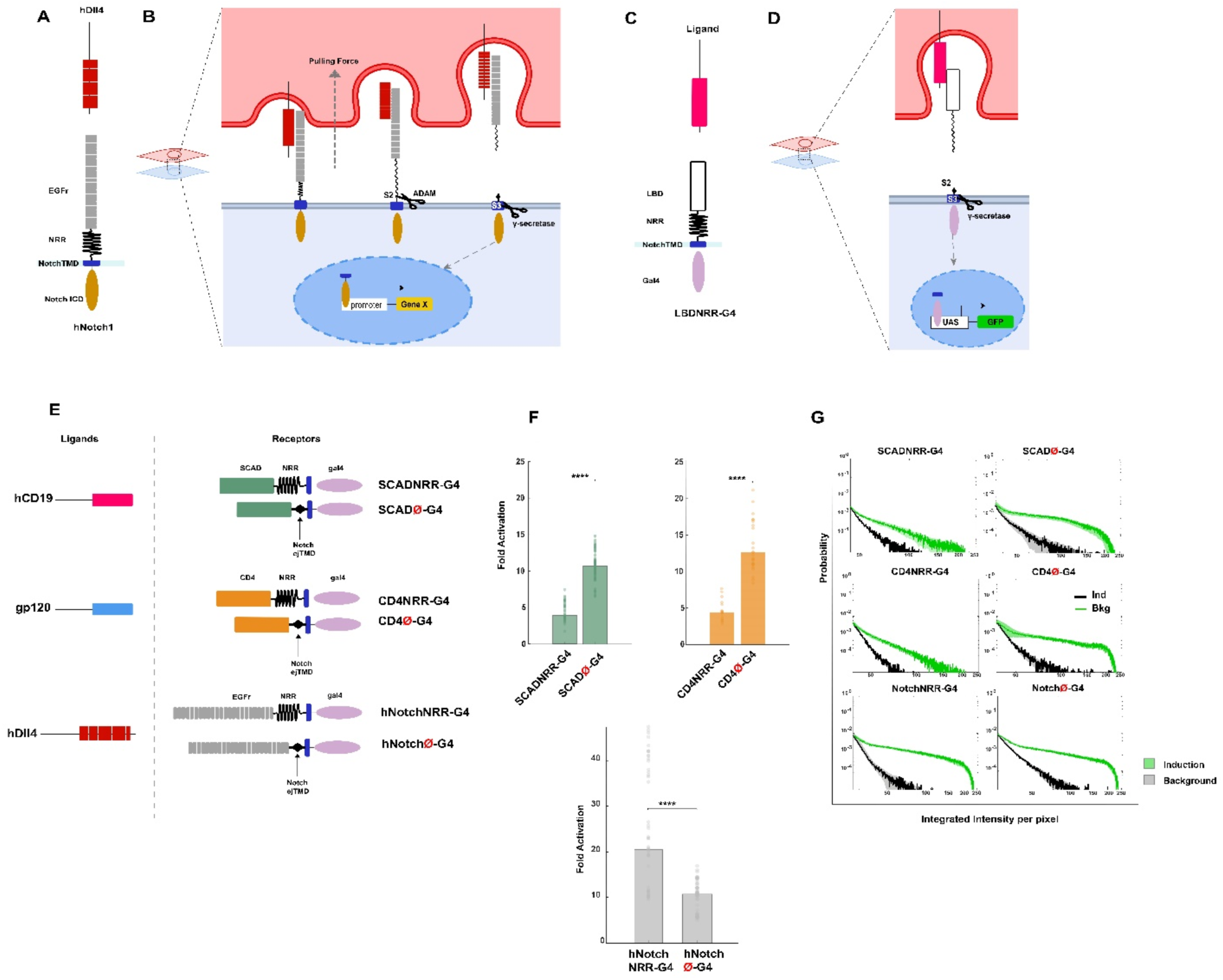
Engineered notch-like receptors withouth a canonical mechanosensing domain are activated by binding to their ligands. **(A)** Diagram illustrating the domains of Notch. **(B)** Interaction of notch and its ligands triggers two sequential cleavages, S2 (within the NRR domain), and S3 (within the TMD), which lead to the release of the intracellular domain (ICD) from the membrane, allowing its translocation to the nucleus where it activates transcription of specific target genes. **(C)** Engineered notch-like receptors LBDNRRG4 contain a ligand binding domain (LBD), the NRR, eJTMD, TMD from human Notch1, and Gal4 in its ICD. The antigen recognized by the LBD is used as a ligand for the activation of LBDNRRG4. **(D)** The mixing of cells carrying LBDNRRG4 receptors with cells expressing the ligand recognized by the LBD release gal4 from the engineered receptor, which translocates to the nucleus and activates transcription of the UAS-gfp reporter. **(E)** SCADNRRG4 receptors contain a single chain antibody domain (SCAD), the NRR, eJTMD, TMD from human Notch1, and Gal4 in its ICD. The human CD19 antigen is recognized by the SCAD (hCAR) domain and is used as a ligand to activate SCADNRRG4. CD4NRRG4 carries the human CD4 extracellular domain in its LBD, and it binds to the gp120 antigen from HIV. SCADØG4 is a variant of SCADNRRG4 in which the NRR domain has been deleted. CD4ØG4 is a variant of CD4NRRG4 missing the NRR. Notch with and without NRR carrying GAL4 as intracellular domain. **(F)** Human notch1 and engineered notch-like receptors carrying SCAD or CD4 can be activated upon binding to their ligands (hDLL4, hCD19, and gp120, respectively) regardless of the presence or absence of the NRR domain. **(G)** Histograms showing normalized probability distribution of intensities above the threshold (median is shown as solid lines and standard deviation shown as thin lines). The green trace is ligand-dependent activation, and the black trace is ligand-independent background.

Recent works have taken advantage of the molecular mechanism of notch activation by using the NRR domain as a mechanoreceptor to monitor cell-cell interactions (Gordon *et al*, 2015; Morsut *et al*, 2016; He *et al*, 2017a; Huang *et al*, 2016). In these designs, the engineered receptors contain an LBD, followed by NNR, the ∼12 aa ejTMD, a transmembrane domain (TMD), and an ICD, which usually contains a transcription factor (**Fig. 1C**). Similarly, to the mechanism of notch activation, in the engineered receptors, binding of the ligand to the LBD of the engineered receptor is thought to trigger a partial unfolding and cleavage in the NRR (S2 site), followed by a second cleavage in the TMD (S3 site) such that an intracellular transcription factor moves to the nucleus and regulates transcription (**Fig. 1D**).

Interestingly, some engineered notch-like receptors lacking the NRR can be activated upon ligand binding, although the mechanism responsible for their activation is not known (Zhu et al, 2022). Here we observed that a ∼12 aminoacid (aa) sequence in the extracellular juxtatransmembrane region (ejTMD) of human Notch1 can act as a mechanoreceptor. In addition, the ∼12 aa ejTMD from notch genes from multiple animal species (chicken, xenopus, zebrafish and *Drosophila*) and other unrelated transmembrane proteins (erbB4, N-CAM, and E-Cadherin) can also act as ligand-dependent mechanosensors. Moreover, multiple artificial sequences of ∼12 amino acids without any specific sequence or apparent structure also act as mechanosensors in engineered notch-like receptors. Finally, we demonstrated that a *Drosophila* Notch gene containing the 12 aa eJTMD but lacking the NRR domain is activated *in vivo* by the physiological forces that occur during early embryonic development. These results indicate that short (∼12 aa), disordered amino acid sequences that are common in many proteins are capable of acting as mechanosensors, suggesting that mechanical force may regulate many more cellular processes than previously suspected.

## Results

### Notch and Notch-like receptors can be activated without a NRR domain

We recently designed a genetically encoded system based on Notch to monitor synaptic interactions in transgenic animals called TRACT (Huang *et al*, 2017). We observed that our engineered notch-like receptors were inefficiently displayed on the plasma membrane. To investigate the possibility that the poor display of engineered receptors was due to the interference between the LBD and NRR in chimeric constructs, we compared the effects of including or omitting the NRR downstream from the LBD. To test this hypothesis we generated two hybrid proteins: (i) SCADNRRG4, whose LBD is a single chain antibody domain (SCAD) that recognizes the human CD19 (Porter *et al*, 2011) (hCD19), the 300 aa of NRR from human Notch1 (hN1), the ejTMD (12 aa) and TMD (24 aa) from hN1 and the yeast transcription factor Gal4 in its ICD, and (ii) SCADØG4, a variant of SCADNRRG4 that does not contain the NRR (**Fig. 1E**). We observed that there were no significant differences in the cell membrane display between SCADØG4 and SCADNRR G4 (data not shown). We observed that cells expressing hCD19 (the ligand that is recognized by the SCAD) activated cells expressing SCADØG4 (the hybrid molecule without NRR) more efficiently than SCADNRRG4 (the hybrid molecule with NRR) (**Figs. 1F and 1G**) (10.68 versus 4.25-fold induction; p=6.5 x 10-5).

Some engineered notch-like receptors lacking the NRR can be activated upon ligand binding, although the mechanisms underlying this activation are not known (Zhu et al, 2022). It has been reported that several large domains (∼100 aa) from multiple proteins can act as mechanosensors (Hayward et al. 2019). Because SCADØG4 did not contain the NRR but could still be activated upon binding to its ligand, we wondered whether the SCAD domain in SCADØG4 could be acting as a ligand-dependent mechanosensor. To test this idea, we generated new hybrid receptors, CD4NRRG4 and CD4ØG4 (containing or lacking the NRR), in which the LBD was switched from SCAD to the ECD from the human CD4 antigen (**Fig. 1E**). We generated stable cell lines expressing CD4NRRG4 and CD4ØG4 and used fluorescent activated cell sorting (FACS) to select for cells with comparable surface expression levels of CD4NRRG4 and CD4ØG4. We placed cells expressing CD4NRRG4 and CD4ØG4 on a plastic surface coated with an antibody that recognizes CD4 and observed that CD4ØG4 had higher levels of induction than CD4NRRG4 (**Fig. 2B**) despite the fact that CD4ØG4 did not have the NRR domain. In addition, we observed that adding the antibody against CD4 into the tissue culture medium failed to activate the receptor, indicating that activation of our engineered receptors requires a mechanical force that cannot be generated by ligands in solution (data not shown). Moreover, we observed that there was no difference between CD4NRRG4 and CD4ØG4 on the plasma membrane display (**Suppl. Fig. 1**). This result indicates that it is possible to obtain force-dependent, ligand-dependent activation of engineered molecules without NRR, and with two different LBDs (CD4 or SCAD) confirming a recent report (Zhu et al, 2022).

**Figure 2.**
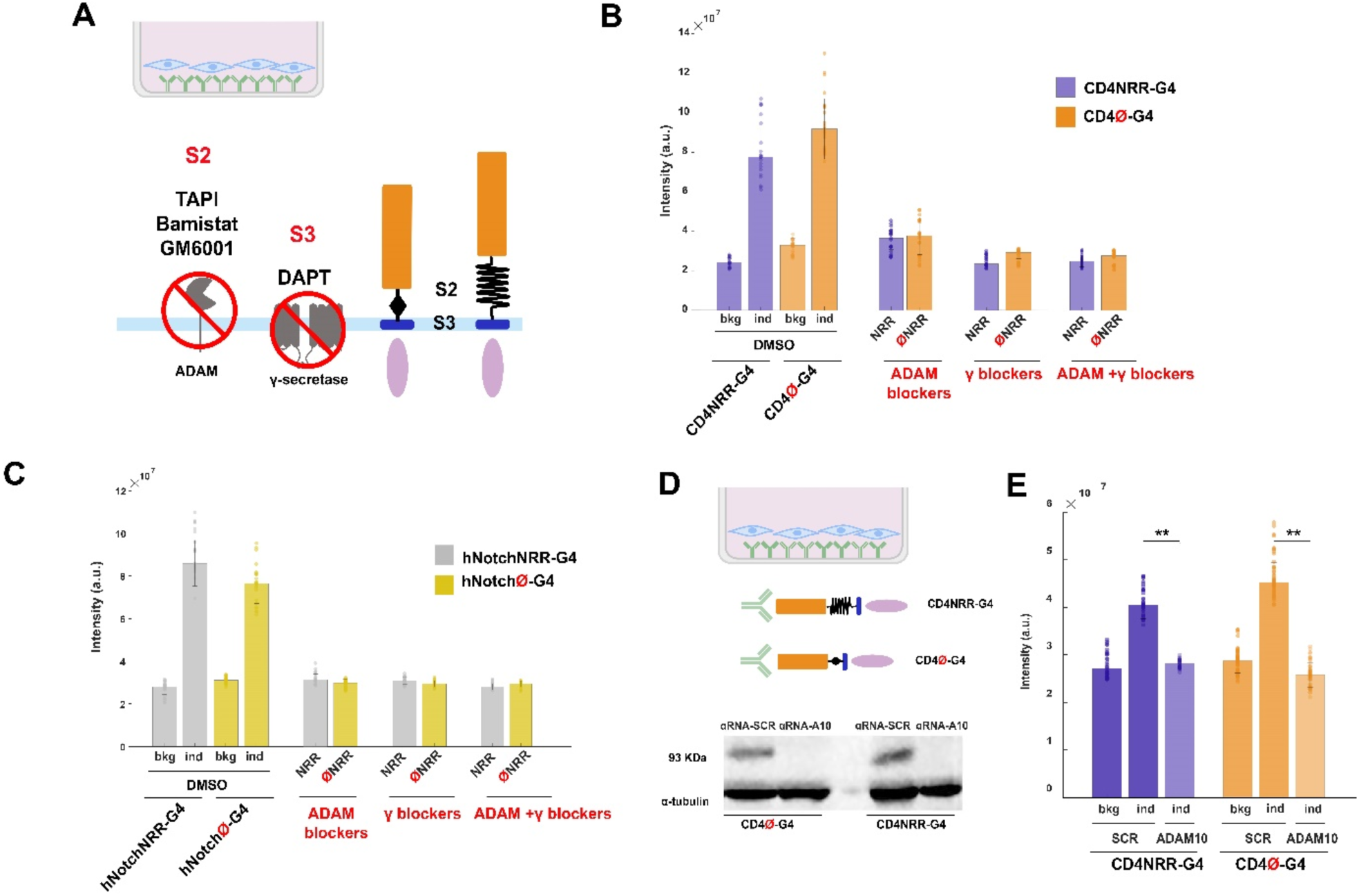
Activation of notch and notch-like receptors withouth a canonical mechanosensing domain requires ADAM protease and gamma-secretase. **(A)** (Top) Schematic representation of induction by immobilized antibody for CD4NRRG4, and CD4ØG4, hNotchNRRG4, and hNotchØG4 receptors. (Bottom) Chemical blockers for ADAM proteases and gamma-secretase inhibit cleavage of S2 site within the NRR, and the S3 site within the TMD, respectively. **(B)** ADAM chemical blockers: TAPI, bamistat and GM6001 reduce the induction of CD4NRRG4 and CD4ØG4. The gamma secretase blocker DAPT reduces activation of CD4NRRG4 and CD4ØG4. **(C)** Decrease of ligand-dependent induction from Notch receptors by ADAM and gamma-secretase blockers. **(D)** Genetic ablation of ADAM 10 blocks activation of receptors with and without NRR. Western blot showing the ablation of ADAM10 protein from receptor cell lines infected with CRISPR-Cas9 and a gRNA against ADAM10 gene or a scramble gRNA (control). α-tubulin is shown as a loading control. **(E)** Genetic deletion of ADAM 10 reduced activation of CD4NRRG4 and CD4ØG4 almost to the same level as the ligand-independent background.

We wondered whether the interaction between the anti-CD4 antibody and the CD4ØG4, or between hCD19 and SCADØG4 could trigger activation of the receptor because the binding of antibody-antigen is essentially irreversible, and it could generate stretching forces that are not normally seen in cell-cell interactions under physiological conditions. To study a ligand-receptor interaction with binding forces observed in cell-cell interactions, we generated cell lines expressing the HIV glycoprotein gp120 (Mao *et al*, 2012), which binds to human CD4 to enable entry of HIV into T-lymphocytes (**Fig. 1E**). We mixed cells expressing gp120 with cells carrying CD4NRRG4 and CD4ØG4 and observed higher activation of receptors lacking the NRR (**Figs. 1F and 1G**, 12.06-fold versus 4.90 fold induction; p=3.69 x 10-8), indicating that receptors without the NRR could also be activated by ligand-receptor pairs that bound to each other reversibly.

We observed that Notch-like engineered receptors carrying a LBD but lacking the NRR could be activated. To investigate whether the NRR was necessary for the ligand-induced activation of Notch itself, we generated constructs with modifications on the ECD of human Notch1 (hN1). The hN1 ECD can be subdivided into three main domains: (i) the LBD, (ii) the NRR, and (iii) the ejTMD (Kopan & Ilagan, 2009). The ligand binding domain consists of 36 EGF repeats. To further examine the role of NRR on notch signaling, we generated two constructs. (i) hNotchNRRG4, encoding the 36 EGF repeats, NRR, ejTMD and TMD of hN1, followed by Gal4, and (ii) hNotchØG4, a variant of hNotchNRRG4 without NRR (**Fig. 1E**). We investigated the inducibility of the modified Notch constructs with a cell line that expressed human delta4 (hDll4), one of the ligands for hN1. We observed that there was comparable activation for hNotchNRRG4 and NotchØG4 constructs regardless of whether they contained the NRR domain or not. (**Figs. 1F and G**; hNotchNRRG4=20.49-fold; NotchØG4=10.80-fold; p=1.49 x 10-7, see **Suppl. Fig. 2A**). Thus, both hNotch and hNotch-like receptors lacking the NRR domain can be efficiently activated upon binding to their cognate ligands.

### Requirements for ligand-dependent cleavage of receptors lacking an NRR

Because the ECDs of SCADØG4, CD4ØG4, and NotchØG4 do not contain the NRR or the canonical S2 site that is usually cleaved upon ligand-receptor interaction in the Notch pathway, we tested additional conditions, examining whether the activation of receptors missing NRR occurred by a mechanism similar to that of wild type notch, requiring ADAM proteases (for an S2-like cleavage), and gamma-secretase (for an S3 cleavage) (**Fig. 2A**). First, we observed that adding to the cell culture medium a cocktail with ADAM protease blockers (batismastat, GM6001 and TAPI), strongly reduced the activation of receptors with or without NRR (**Figs. 2B and C**) - ( reduction of induction by CD4NRRG4 and CD4ØG4 by 81.30% and 93.34%; respectively;, p=3.50 x 10-4)., and NotchNRRG4 and NotchØG4 by 91.72% and 104.48%, respectively ; p=4.46 x 10-8 - (note - addition of the ADAM blockers reduced ligand-dependent induction below ligand-independent background levels without blockers, accounting for the reduction of induction greater than 100% in some cases). This suggests that even in the absence of NRR, or a canonical S2 site, the binding of ligand to CD4ØG4 and NotchØG4 triggers a cleavage mediated by ADAM metalloproteases. Because ADAM protease blockers are not strictly specific for any individual member of the ADAM metalloproteases, we generated cell lines in which the ADAM10 gene (the key ADAM member involved in ligand-dependent Notch activation) was mutated by CRISPR (**Fig. 2D**). We observed that induction of CD4NRRG4 or CD4ØG4 was strongly reduced in ADAM10-deficient cell lines (**Fig. 2E**) (reduction of induction of CD4NRRG4 and CD4ØG4 by 80.84% and 115.86%, respectively (p=0.0072). These observations indicate that although there is no NRR or canonical S2 site in the CD4ØG4 or hNotchØG4 receptors, the interaction of these receptors with their ligands trigger an ADAM-dependent cleavage similar to that observed when wild-type notch binds to its ligands. Second, the gamma-secretase blocker DAPT abolished the activation of the receptors CD4NRRG4, CD4ØG4, hNotchNRRG4, and hNotchØG4 (**Figs. 2B and C**) (reduction by 98.15% and 107.76% for CD4NRRG4, CD4ØG4, respectively; p=7.31 x 10-8), and reduction by 93.29% and 102.76% for hNotchNRRG4 and hNotchØG4, respectively; p=4.17 x 10-7) indicating that the activation of the receptors with or without NRR also required intramembrane proteolysis mediated by gamma-secretase.

These experiments indicate that both for Notch (hNotchØG4) and engineered receptors (SCADØG4 or CD4ØG4), ligand binding induced cleavage of the receptor in the absence of the NRR domain or a canonical S2 site that could be inhibited by ADAM blockers. This observation indicated that these constructs may contain sites with the potential to be cleaved by ADAM proteases upon ligand binding to the different LBDs. The eJTMD that were present in these constructs included 12 aa (the sequence QSETVEPPPPAQ) from the hN1 eJTMD starting after the canonical S2 site (Ala/Val) of hN1 until the first amino acid of the hN1 TMD (Leu). To investigate whether the activation of the engineered receptors was specifically dependent on the hN1 eJTMD we generated constructs with different types of eJTMDs. To this end, we generated constructs in which the eJTMD from chicken notch1, xenopus notch, zebrafish notch1, human notch2, and *Drosophila* notch were placed between CD4 and hN1 TMD (**Fig. 3A**). We observed in all these cases that eJTMDs from notch genes from different species were sufficient to induce ligand-triggered cleavage of the engineered receptors in the absence of NRR (**Fig. 3A**; hNotch1=11.63-fold, hNotch2=4.16-fold, p(hN2)=3.85 x 10-10; chNotch=7.43-fold, p(ch)=1.65 x 10-7; zfNotch=7.97-fold, p(zf)=5.10 x 10-6; xNotch=7.48-fold, p(x)=8.10 x 10-7; dNotch=3.77-fold, p(d)=7.35 x10-10). These experiments indicate that short sequences (∼12aa) in the eJTMD of different notch molecules can act as ligand-dependent mechanosensors when placed in heterologous molecules.

**Figure 3:**
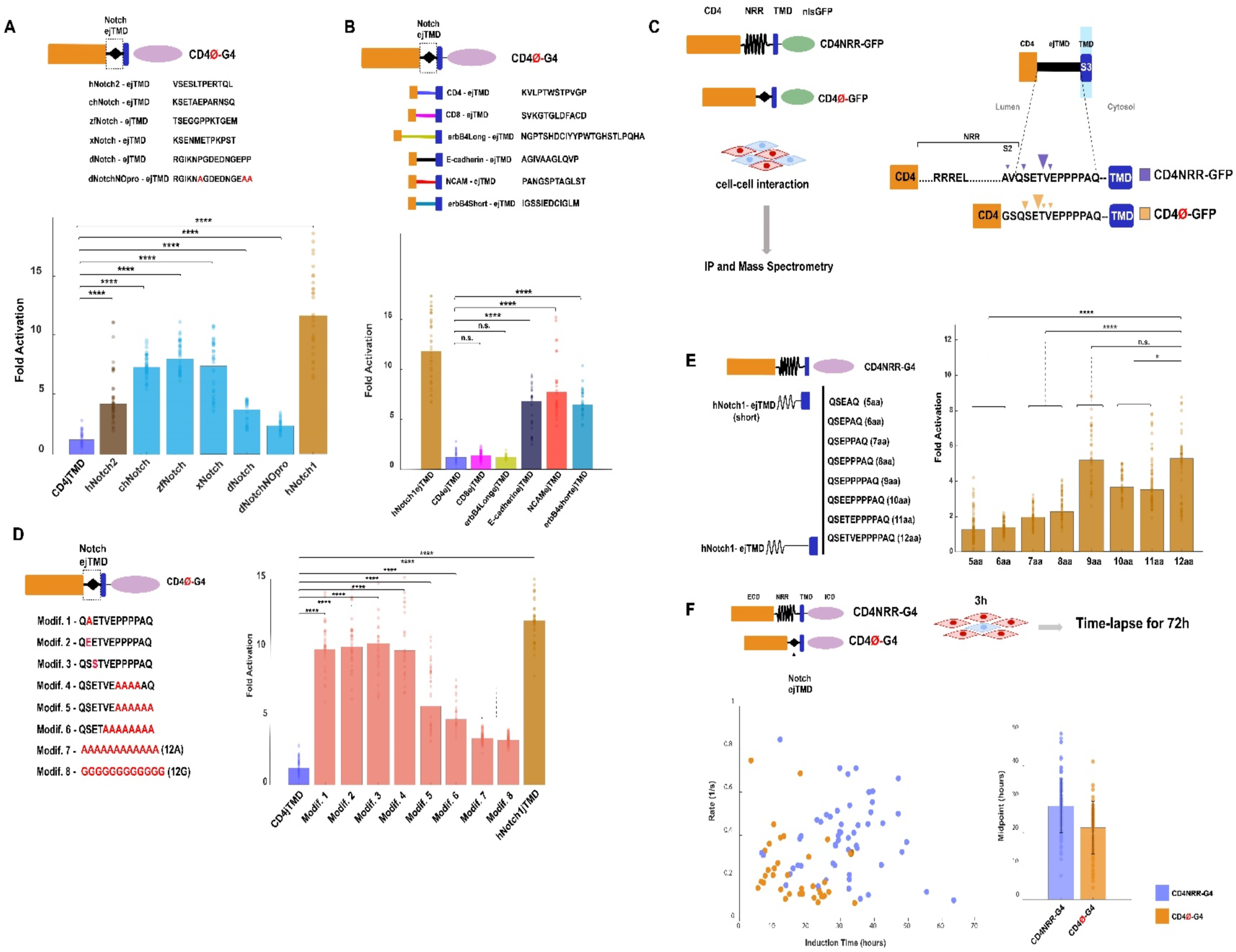
Extracellular juxtatransmembrane domains from different membrane proteins and without defined structure act as a mechanosensor that respond to cellular forces. **(A)** The eJTMD from notch molecules derived from multiple species produce ligand-dependent activation of notch-like receptors. (Top) sequences of eJTMD from Notch molecules from different species. (Bottom) Fold Activation showed inducibility of eJTMD from notch molecules derived from multiple species. **(B)** The eJTMD from the human CD4, CD8 antigens, or the long isoform from the erbB4 receptor do not lead to ligand-dependent activation of Notch-like receptors, whereas the eJTMD from E-caherin, NCAM, and short isoform of erbB4 produce ligand-dependent activation of notch-like receptors (fold activation). **(C)** Left: Diagrams showing the experimental strategy to biochemically identify the cleavage sites in receptors CD4NRR-GFP and CD4Ø-GFP. Right: Notch-like receptors with NRR are preferentially cleaved in a site in the the eJTMD between threonine (T) and Valine (V), and those without NRR get preferentially cleaved in the eJTMD between glutamic acid (E) and threonine (T). **(D)** Multiple changes in the aminoacid sequence of the human Notch1 eJTMD (left) do not block its ability to trigger ligand-induced activity of notch-like receptors (right). **(E)** Ligand-dependent activation of notch-like receptors with NRR depends on the length of the eJTMD; (left) sequences of eJTMDs of different lengths and (right) fold activation induced by ligand-receptor binding with eJTMDS of different lengths. **(F)** (Left) Scatter plot showing the rate at which the cells get induced vs the induction time, and (right) time to reach half-maximum ligand-dependent activation for notch-like receptors with and without NRR domains.

To investigate whether the eJTMD from other molecules unrelated to notch could also act as mechanosensors, we generated constructs encoding 12 aa of the eJTMD from several different type I transmembrane proteins involved in cell-cell signaling or adhesion: NCAM, ECadherin, erbB4 (short and long isoforms), CD4, and CD8 (**Fig. 3B**). We observed that whereas receptors carrying the eJTMDs from ECadherin, NCAM, and short isoform of erbB4 (erbB4S) produced ligand-dependent activation of notch-like receptors (6.49, 8.24, and 6.43 fold induction, respectively), those carrying eJTMDs from CD4, CD8, and long isoform of erbB4 (erbB4L) did not (1.09, 1.40 and 1.68 fold activation, respectively). Although all these constructs carried the same TMD (from hN1), only some eJTMDs conferred ligand-dependent activation. To test whether the hN1 TMD was necessary for the observed ligand-dependent activation in the absence of an NRR domain, we generated receptors carrying the CD4 ECD, the hN1 eJTMD and the TMDs from three type-I transmembrane domains (sevenless, torso, an ß-APP) followed by esn. We observed that receptors carrying the hN1 eJTMD and these 3 different TMDS could be activated as those carrying the hN1 TMD (**Suppl. Fig. 3**). This result indicates that the hN1 TMD is neither necessary not sufficient for the ligand-dependent activation of the engineered receptors lacking an NRR, and that short (∼12 aa) eJTMD sequences from molecules other than notch can act as mechanosensing domains that can be grafted into hybrid molecules to render them ligand-inducible.

To biochemically identify the putative cleavage sites in the eJTMD in our receptors, we generated two receptors, CD4NRRGFP or CD4ØGFP, whose ICD contained GFP, which was used as a tag for purification by immunoprecipitation with antiGFP nanobodies. Cells expressing CD4NRR-GFP or CD4Ø-GFP were mixed with cells expressing gp120 (the CD4 ligand), and the receptors were immunoprecipitated and subjected to mass spectrometry to identify the putative cleavage sites in the eJTMD (**Fig. 3C**). Consistent with previous works, we observed a cleavage site within the NRR in CD4NRRGFP between Ala and Val (A/V), the so-called S2 site (**Fig. 3C**). However, we identified that the most abundant cleavage site was located within the eJTMD (downstream from the NRR) of CD4NRRGFP between T/V. In addition, we also identified two additional cleavage sites within the eJTMD, between Q/S, and V/E. CD4ØGFP is missing the NRR and the canonical A/V S2 site, and it was cleaved in several sites within the eJTMD, with the most abundant site between E/T, followed by Q/S, T/V and V/E. This experiment indicates that receptors missing the NRR (and the A/V S2 site), are still activated by a similar ADAM-mediate proteolytic cleavage in response to ligand binding, and that eJTMD can act as a cellular mechanosensor.

Several constructs missing the NRR but including 12 aa eJTMDs from several different proteins (hN1, notch from different species, NCAM, erbB4S, and E-cadherin) still showed induction upon ligand binding (**Figs. 1F, 3A and B**). However, not all eJTMDs (eg: eJTMD from CD4, CD8, or erbB4 long isoform) could act as ligand-dependent regulators of notch-like receptors (**Fig. 3B**). Cursory inspection of the sequence of these different eJTMDs does not reveal any obvious motif or pattern that hints at which sequences make them susceptible or resistant to cleavage (**Suppl. Fig. 7**). To further investigate whether there were critical aa sequences in the eJTMDs necessary for mechanotransduction, we generated several constructs with the following modifications of the hN1 eJTMD. (i) Q*<u>A</u>*ETVEPPPPAQ (change from S to A), Q*<u>E</u>*ETVEPPPPAQ (change from S to E), (iii) QSSTVEPPPPAQ (change from SE to SS), (iv) and QSET*<u>AAAAAAAA</u>* (change from VEPPPAQ to AAAAAAA). None of these modifications in the sequence of the eJTMD significantly impaired the ligand-induced activation of the receptors (**Fig. 3D**; modif. 1(QAETVEPPPPAQ) 9.95 fold, modif. 2(QEETVEPPPPAQ) 10.16-fold, modif. 3(QSSTVEPPPPAQ) 10.33-fold, and modif. 4 (QSETVEAAAAAQ)-9.62 fold. Changing the last 5 (QSETVEAAAAAAA)) or 7 ((QSETAAAAAAAA)aa of the eJTMD for Ala, still resulted in significant induction by ligand binding (modif. 4 (QSETVEAAAAAA)=5.80-fold;) modif. 5 (QSETAAAAAAAA)=4.77-fold).

Finally, to test the hypothesis that even sequences without any secondary structure could act as mechanoreceptors, we generated constructs in which the eJTMD consisted of 12 ala or 12 gly, two sequences which do not possess any defined structure. Interestingly, even these eJTMD exhibited ligand-triggered activation, albeit significantly weaker than that observed with the Notch1 eJTMD (∼3 fold versus 13 fold, **Fig. 3D**). (**Suppl. Fig. 5C**; p-values between induction and background: hNotch1eJTMD=6.51 x 10-12; modif. 1=1.55 x 10-6; modif. 2=3.02 x 10-12; modif. 3=1.66 x 10-6; modif. 4=1.64 x 10-5; modif. 5=1.24×10-5; modif. 6=6.60 x 10-14; modif. 7=3.23 x 10-17; modif. 8=1.62 x 10-20).

Our experiments showed that not all eJTMD were efficient as mechanosensors, with the human Notch1 eJTMD being the most efficient (∼13 fold induction), and the eJTMD from CD4 and CD8 being inactive. One can imagine several scenarios that could explain the inability of the CD4 and CD8 eJTMD to act as a mechanosensor. For example, the sequence of the CD4 eJTMD could not be cleaved by proteases after ligand binding. Alternatively, this eJTMD could actively inhibit the cleavage by proteases into other potential sites. To distinguish between these two scenarios, we generated constructs that contained the LBD of CD4 and the complete hN1 NRR (including the canonical A/V S2 site) followed by: (i) hN1 eJTMD (12 aa), (ii) the CD4 eJTMD (12 aa), and (iii) a double eJTMD consisting of the the hN1 eJTMD (12 aa) followed by the CD4 eJTMD (12 aa). We observed that although constructs carrying the CD4 eJTMD by itself could not be induced (**Fig. 3B**), the presence of the CD4eJTMD did not inhibit the ability of the hN1 NRR or hN1 eJTMD to be cleaved (**Fig. Supp 4**). Unexpectedly this experiment also indicates that the NRR and/or the hN1 eJTMD can be cleaved when placed farther away from the cell membrane than in their original location (24 aa versus 12 aa).

Our data indicates that multiple sequences of around 12 aa can act as mechanosensors in eJTMD, without requiring the presence of NRR. Can the NRR function as a mechanoreceptor in the absence of a full-length (12 aa) ejTMTD? To test this hypothesis we generated constructs that consisted of the CD4 LBD and the hN1 NRR separated from the TMD by variants of the hN1 eJTMDS of different lengths, ranging from 5 to 12 aa [5 (QSEAQ), 6 (QSEPAQ, 7 (QSEPPAQ), 8 (QSEPPPAQ), 9 (QSEPPPPAQ), 10 (QSEEPPPPAQ), 11 (QSETEPPPPAQ, and 12 (QSETVEPPPPAQ [the wt notch eJTMD]. We observed that shortened eJTMDs with only 5 and 6 aa prevented the activation of the receptor despite the presence of a complete NRR (including the A/V S2 site). In contrast eJTMDs with 7 8 9 10 11 aa had ligand-dependent inductions comparable to those observed with the wt Notch1 eJTMD (12 aa). (Fig 3E (5aa=1.24-fold; 6aa=1.38-fold; 7aa=1.97-fold; 8aa=2.30-fold; 9aa=5.05-fold; 10aa=3.64-fold; 11aa=3.47-fold; and 12aa=5.20-fold). This experiment indicates that for the NRR to act as a mechanoreceptor it requires the presence of an eJTMD, although we do not know if this requirement depends on a specific sequence of aa in the eJTMD, or simply a minimal length of separation between the NRR and the cell membrane.

Our experiments indicate that multiple short aa sequences can act as ligand-dependent mechanosensors and that the NRR is dispensable for the activity of Notch confirming a previous observation (Zhu et al, 2022). Thus, if the NRR is not necessary for activation of Notch this raises the question of what its function is. We observed that although the ligand-dependent activation of CD4ØG4 was higher than that of CD4NRRG4 (∼13 fold versus ∼4 fold), the ligand-independent background signaling of CD4ØG4 was approximately ∼2 fold higher than CD4NRRG4 (**Fig. Sup 6 or Fig. Sup 2A**). We also observed that in CD4NRRG4 and hNotchNRRG4 the presence of the NRR domain also reduced the ligand-independent background, compared to constructs missing it (SCADØG4, hNotchØG4) (**Fig. Sup 2A**). This indicates that one of the functions of the NRR could be to reduce the ligand-independent cleavage of the eJTMD by ADAM proteases. To investigate other potential functions of the NRR domain we measured the temporal dynamics of ligand-dependent activation of CD4NRRG4 and CD4ØG4. To this end, we performed time-lapse microscopy of CD4NRRG4 and CD4ØG4 cells mixed with cells expressing gp120 (the ligand for CD4). We observed that cells with CD4ØG4 reached half-maximal induction significantly faster than those with CD4NRRG4 (**Fig. 3F**) (21.46 ± 7.84 hours versus 27.95 ± 8.08 hours for CD4ØNRRG4 and CD4NRRG4, respectively; p=7.20 x 10-10), indicating that receptor without NRR could accelerate the action of ADAM proteases upon ligand binding.

### Mechanosensing in the absence of NRR in the context of a developing animal

To investigate whether short eJTMDs could act as mechanosensors under physiological conditions in animals, we generated transgenic *Drosophila* carrying two Notch transgenes whose only difference was the presence or absence of the NRR domain. Each of these transgenic flies carry a different construct: (a) dNotchNRR-Gal4, consisting of the full-length *Drosophila* Notch (dNotch) LBD, NRR, eJTMD and TMD followed by Gal4 in its ICD, and (b) dNotchØ-Gal4, a similar construct missing the NRR domain (**Fig. 4A**). The transgenes also included a V5 tag to enable immunodetection of the transgene. Both these constructs were placed under the PREX heat shock promoter, to enable temporal control of transcription, and were inserted into the attP P2 site, to achieve comparable levels of expression between both constructs. To obtain a readout of the activation of the dNotchNRRG4 and dNotchØG4 transgenes, we crossed these flies with transgenic flies carrying a UAS-nlsGFP reporter (**Fig. 4B**). We obtained embryos from these genetic crosses and delivered a heat shock (2 hours long) to embryos within an hour after deposition. We performed immunostaining against the V5 tag to detect the presence of the receptor transgenes and observed that both dNotchNRRG4 and dNotchØG4 transgenes were ubiquitously expressed throughout the embryo in an apparently homogeneous pattern, consistent with the activity of the PREX promoter (**Fig. 4C**).

**Figure 4:**
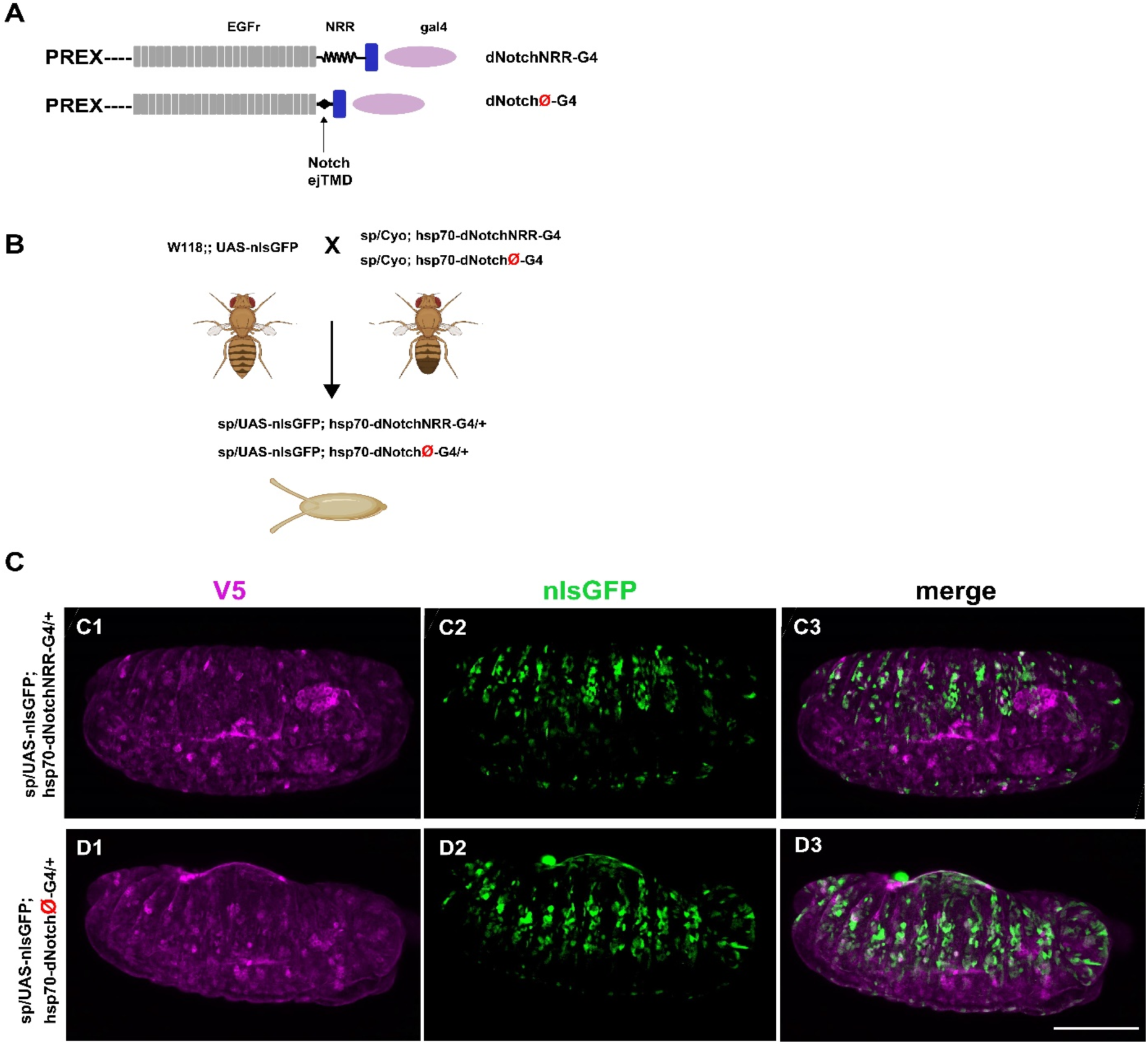
Notch receptors with a short juxtatransmembrane domain are activated by physiological signals in *Drosophila* developing embryos. **(A)** Diagrams of Notch transgenes with (PREX-dNotchNRR-Gal4) and without (PREX-dNotchØ-Gal4) NRR domains. Transgenes were cloned downstream from the Heat-shock promoter PREX. **(B)** PREX-dNotchNRR-Gal4 and PREX-dNotchØ-Gal4 flies were crossed to flies carrying the reporter UAS-nlsGFP to image the activation of GFP expression in ∼10-11 hours old embryos. **(C)** C1 and D1 (purple) Immunostaining against V5 tag reveals the ubiquitous presence of the dNotchNRRG4 and dNotchØG4 receptors, respectively, in 10-11 hours embryos (stage 14); C2 and D2-pattern of expression of GFP from the UAS-nls GFP reporter triggered by activation of the dNotchNRRG4 and dNotchØG4 receptors in embryos; C3-D3 – merge of V5 and nlsGFP expression pattern. The pattern of activation of Notch as revealed by the UAS-nlsGFP reporter is similar regardless of the presence or absence of the NRR domain in the transgenes. Scale bar in D3 is 100 µm and applies to C1-D3.

## Discussion

Mechanical forces regulate many biological processes, and multiple proteins have specialized mechanosensing domains to respond to those forces. Two of the first mechanosensing domains identified were the A2 domain from the von Willebrand factor (VWF), and the NRR domain from Notch. The VWF A2 domain is a 193 aa region from the von Willebrand factor (VWF). In response to hemodynamic stresses in the bloodstream the A2 domain is stretched, and it exposes a site that can be cleaved by extracellular proteases (Zhang *et al*, 2009a). A similar mechanism regulates the activity of Notch; whereby mechanical forces associated with ligand binding induce exposure of a site that is then cleaved by extracellular proteases. It is currently believed that the notch NRR, a sequence of around 300 amino acids between the EGF repeats and the TMD, is involved in the ligand-dependent activation of notch. This belief is based on several lines of evidence. First, the NRR cleavage site has homology to the Sea Urchin enterokinase agrin-like (SEA-like) domain, a target for ADAM 10/17 proteases (Hayward *et al*, 2019). Second, crystallographic studies indicate that the so-called S2 cleavage site is buried inside of NRR, and thus protected from ADAM metalloproteases (Gordon *et al*, 2009). It is thought that ligand binding to notch stretches the NRR and exposes the S2 site such that it can then be cleaved by the ADAM proteases. Third, some leukemias are caused by mutations in the NRR that lead to ligand-independent activation of notch molecules. It is thought that these mutations cause a misfolding of the NRR such that it does not occlude the S2 site (Weng *et al*, 2004), and thus it can be accessed by the ADAM proteases even in the absence of ligand binding. Fourth, adding the NRR into different artificial receptors enables the ligand-dependent of the hybrid molecules upon ligand binding (Gordon *et al*, 2015; Morsut *et al*, 2016; He *et al*, 2017b; Huang *et al*, 2016; Huang *et al*, 2017). Thus, it is thought that NRR is the critical element to impart mechanosensing capabilities into these hybrid proteins, and that NRR is a specialized protein domain that acts as a portable ligand-dependent mechanosensor.

Interestingly, some engineered notch-like receptors lacking the NRR can be activated upon ligand binding, although the mechanisms underlying this activation are not known. (Zhu et al., 2022). Here we demonstrate that a ∼12 amino acid sequence located downstream from the NRR, and immediately upstream of the transmembrane domain (the extracellular juxtatransmembrane domain or eJTMD) of notch is sufficient to achieve ligand-dependent cleavage in response to physiological mechanical forces. Moreover, our data demonstrates that there is no strict requirement in the sequence of the eJTMD to enable ligand-dependent mechanosensing, as we have shown that multiple changes in several aminoacids from the hN1 eJTMD still enable ligand-dependent cleavage in response to force. In addition, we also demonstrate that ∼12 aa eJTMD from other notch molecules (chicken, fish, frog, drosophila) and from proteins other such as E-cadherin, erbB4S, and NCAM that have not been previously reported to be regulated by mechanical forces can also turn proteins into mechanosensors. Even short (around 12 aa) sequences of amino acids without defined secondary structure can also turn different proteins into ligand-dependent mechanosensors when placed in the eJTMD, albeit with significantly reduced efficiency. However, not all sequences can act as mechanosensors and we observed that some eJTMDs, such as the eJMTDs from CD4 or CD8, do not enable ligand-mediated activation of notch-like receptors. Finally, we have observed that a Notch protein missing the NRR but carrying the 12 aa of its eJTMD can be activated *in vivo* by the physiological relevant mechanical forces that occur in the developing *Drosophila* embryo and produce a pattern of activation comparable to that of the wild-type Notch.

The results from our experiments indicate that multiple short aa sequences can act as ligand dependent mechanosensors and that a Notch receptor without the NRR domain can be activated upon binding to its natural ligands *in vitro* and *in vivo*. This raises the question of which is the function of NRR, a large protein domain (∼300 aa) that appears to be conserved in Notch molecules across all metazoans. We observed that although the ligand-dependent activation of CD4ØG4 was around 3 fold higher than CD4NRRG4, the ligand independent activation of CD4ØG4 was approximately two folds higher than CD4NRRG4, suggesting that NRR domain play an essential role in tightening the cleavage of the S2 site and preventing non-specific, ligand independent activation. Moreover, our results from time-lapse indicate that the engineered receptor with NRR presented a slower dynamic than the CD4ØG4receptor, suggesting that the NRR could facilitate the interaction between the ADAMs and S2 cleavage site, allowing a more efficient cleavage by these proteases. Another possibility is that after the mechanical tension has been exerted by the interaction with the ligand with the receptor, the NRR maintains the protein in an “open” state increasing the length of time that ADAM proteases can cleave the exposed S2 site. However, our *in vivo* results did not show any gross differences in the *Drosophila* embryo’s activation pattern carrying the Notch receptor with and without NRR. Future experiments will be needed to evaluate how Notch receptor dynamics is affected by the NRR domain *in vivo*.

These observations indicate that adding an appropriate eJTMD can convert different kinds of type I transmembrane proteins that do not have any specific mechanosensing domains into mechanosensors that respond to physiological forces caused by cell-cell interactions. These proteins have different ligand-binding domains upstream of the eJTMDs. We have tested domains including human CD4, human and mouse CD19, single chain antibody domains that recognize mouse and human CD19, and human and Drosophila Notch (without their NRR). One can envision two likely scenarios that could account for these results. In one model, the eJTMDs that we have identified are actual mechanosensors by themselves such that the mechanical tension caused by ligand binding stretches the eJTMDs and changes their secondary structure, making them more susceptible to ADAM cleavage. This model predicts that many short, unstructured domains can act by themselves as mechanoreceptors and is reminiscent of the requirements for activation domains in transcription factors. Originally it was thought that these transcription activation domains would need to have precise, defined structures to interact with RNA polymerase II. However, it was later found that many short, unstructured aa sequences could act as potent transcription activators. For instance, it was shown that different peptides sequences carrying an excess of acidic residues worked as activation domains when they are attached to DNA binding domains (Ma & Ptashne, 1987). In a second model, the mechanical forces caused by ligand binding do not change the structure of the eJTMD, but they simply displace some other domains of the protein that may be occluding it. The displacement of these other domains away from the eJTMD would increase their accessibility to the ADAM proteases, inducing ligand-dependent cleavage. The eJTMD is located immediately on top of the cell membrane, so it seems likely that the ADAM proteases may be limited in their access to the eJTMD. In this scenario all the different ligand-binding domains that we have tested in our experiments would fold in such a way that they protect the eJTMDs from cleavage by ADAM proteases. This model would suggest that ligand binding may significantly distort the secondary structure of many proteins. Investigating these scenarios will require biophysical experiments aimed at measuring whether physiological mechanical forces are sufficient to change the structure of these short peptide sequences in the eJTMDs, or to cause displacement of different protein domains that may be protecting the eJTMD.

The observations reported here indicate that mechanosensing in proteins does not require large, specialized domains, that mechanosensor proteins may be quite common, and suggest that mechanical force sensing may regulate many more cellular processes than previously suspected.

## METHODS

### Gene constructs

All constructs were generated using standard restriction enzyme-based and Gibson cloning (Gibson et al., 2008). ID3NRRG4 has been described previously (Huang et al., 2016). All constructs for in vitro experiments were subcloned into the FUWG lentiviral backbone (Lois et al., 2002).

SCADNRRG4 was constructed by fusing a single-chain antibody (SCAD) that recognizes the human CD19, the NRR, and TMD from human Notch1 and esn (a variant of Gal4) as the intracellular domain. The NRR domain and TMD comprised amino acid 1446-1880 of human Notch1. The entire SACDNRRG4 was subcloned into the FUWG lentiviral backbone (Lois et al., 2002). All the receptors were cloned following the same logic, introducing different modifications described in the figures or Supplementary Tables. All the gene blocks were generated by Integrated DNATechnologies (IDT).

The SCADNRResn constructs, and its variants, contain a SCAD that recognizes the human CD19. The ligand for the SCAD-carrying receptors is the human CD19 cloned into FUW.

The CD4NRResn constructs, and its variants, contain the extracellular domain of human CD4. The ligand for the CD4-carrying receptors is the HIV antigen gp120 cloned into Moloney retroviral vector.

The human Notch constructs (1-36NRRG4 and 1-36ØG4) contain the ECD (with or without NRR) and TMD from human Notch1, followed by esn.

The Drosophila Notch constructs (PREX-dNotchNRR-Gal4 and PREX-dNotchØ-Gal4) contain the ECD (with or without NRR) and TMD from Drosophila Notch1 followed by esn. The inserts dNotchNRR-Gal4 and dNotchØ-Gal4 were cloned under the heat shock promoter PREX and integrated into the attp P2 site.

For those receptors with different length of the eJTMD carrying NRR, gene block inserts (5 aa, 6aa, 7aa, 8aa, 9aa,10aa, 11aa, 12aa) were subcloned into CD4Øesn.

### Cell culture

CHO cells and their derivatives were grown on tissue-culture grade plastic plates in MEM (10-010-CV, CorningCellgro), supplemented with 10% FBS, and 1% penicillin, streptomycin, and L-glutamine.

HEK293 cells, HeLa cells, and their derivatives cell lines were grown in DMEM, supplemented with 10% FBS, 1%penicillin, streptomycin, and L-glutamine, and 1% sodium pyruvate.

All cells were grown at 37°C in 5% CO2 in a humidified atmosphere. Cells were passed every 1-2 days, depending on confluency, using 0.05% Trypsin-EDTA.

### Lentivirus production

All the lentiviruses were produced by transfection on 293 HEK cells as described in (Lois et al., 2002).

### Generation of Stable Cell lines

Stable cell lines were produced and stored as previously described (Lois et al., 2002), using the plasmids described above. The cells were infected with the specific lentiviral vector. To generate stable lines of emitter cells, CHO-K1 cells were infected by retrovirus expressing either mCD19mCherry, hCD19tdTomato or gp120mCherry. To generate UAS-GFP reporter HeLa cell lines (used for CRISPR-ADAM10 experiments), HeLa cells were infected with a UAS-GFP lentivirus. UAS-GFP reporter HeLa cells were sorted using a FACS device based on GFP intensity to generate homogeneous stable cell lines.

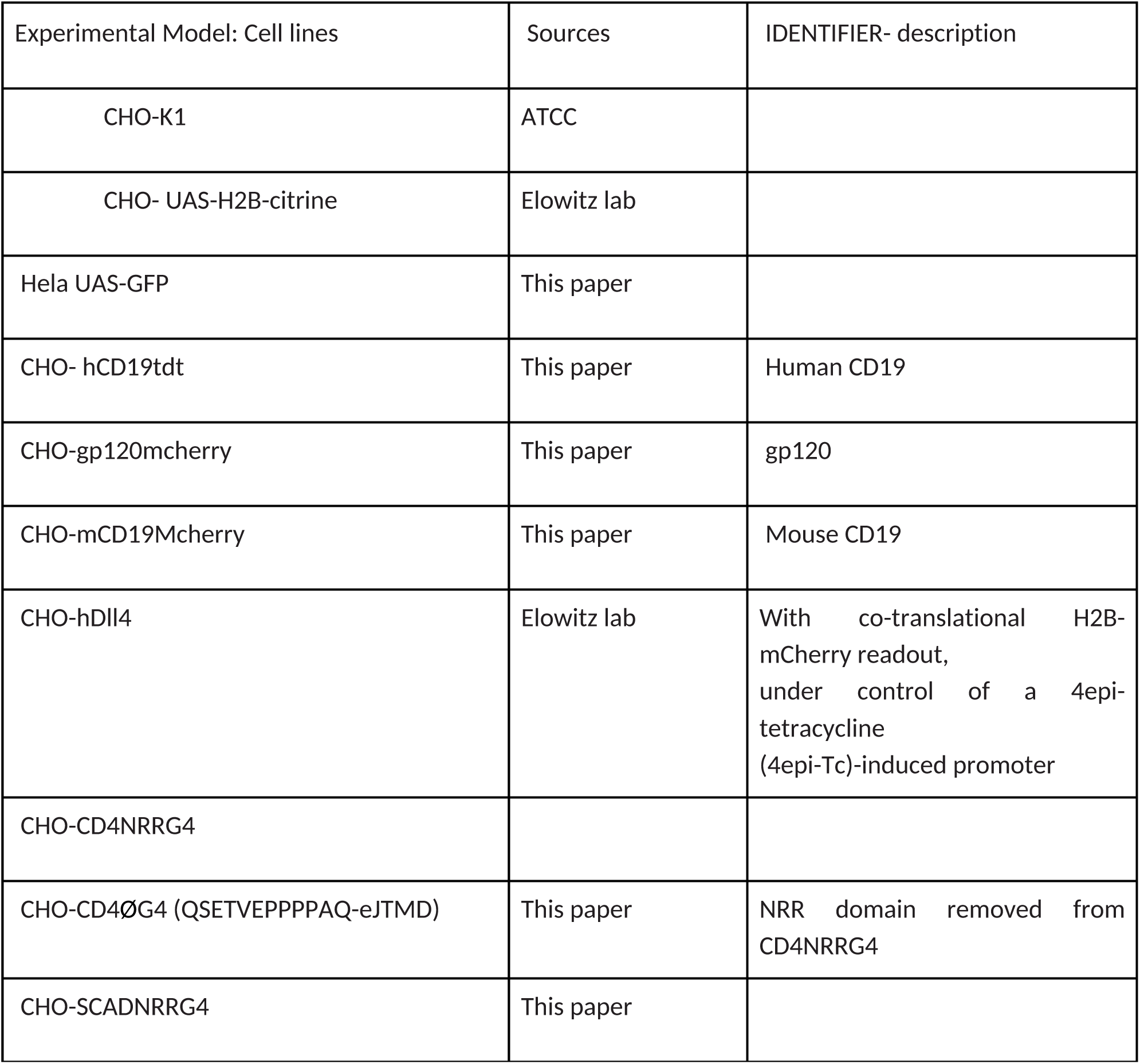
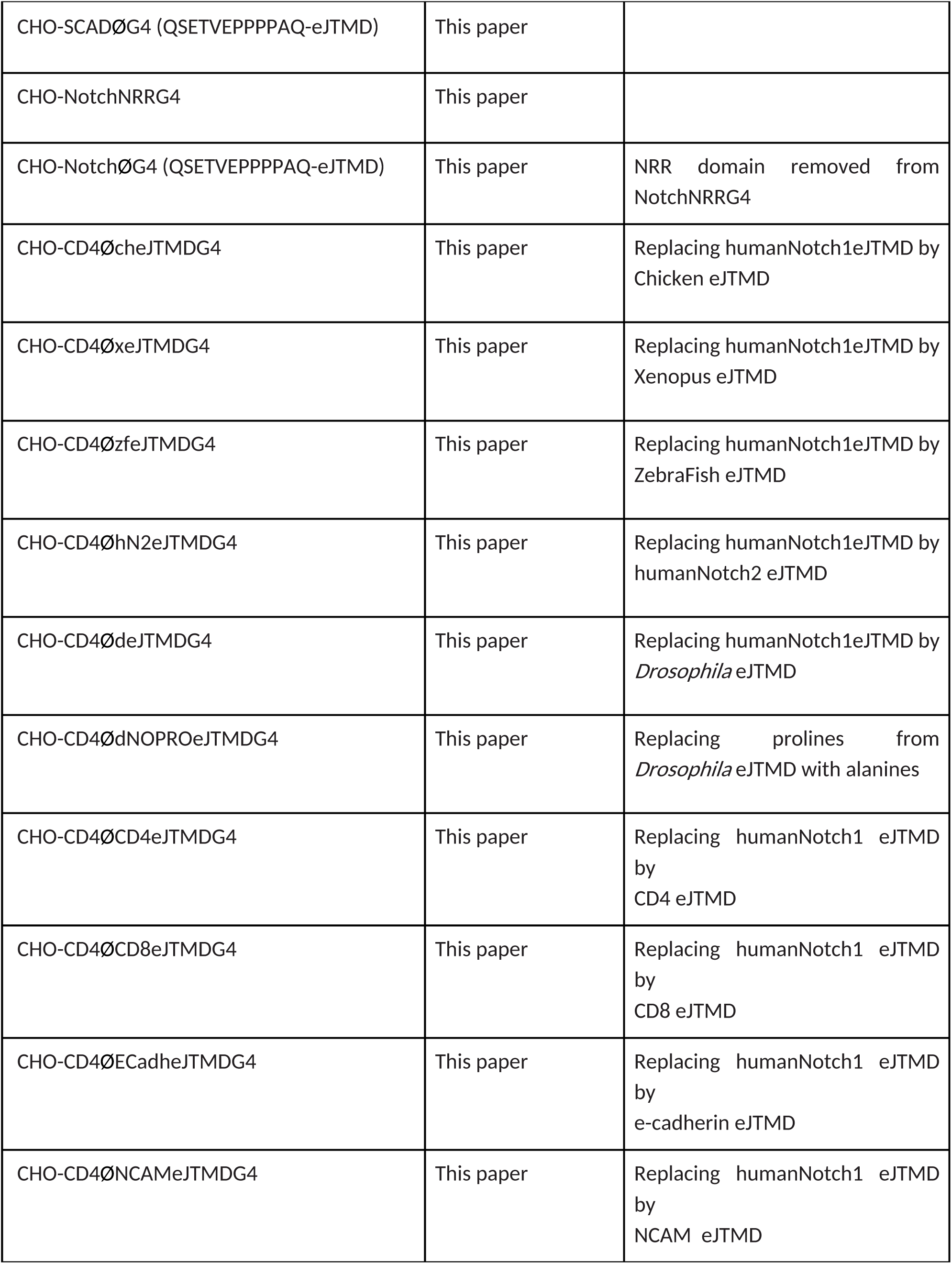
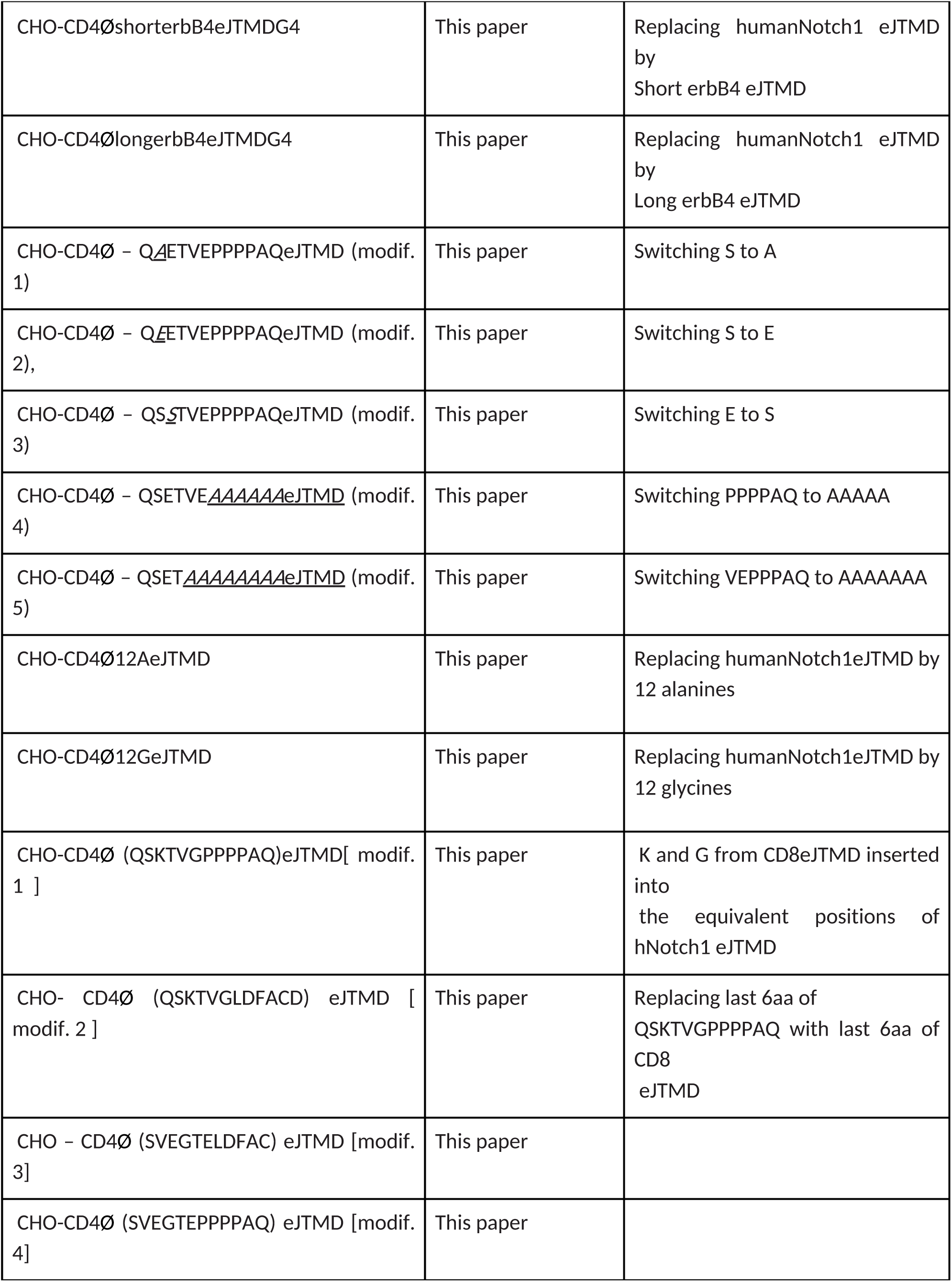
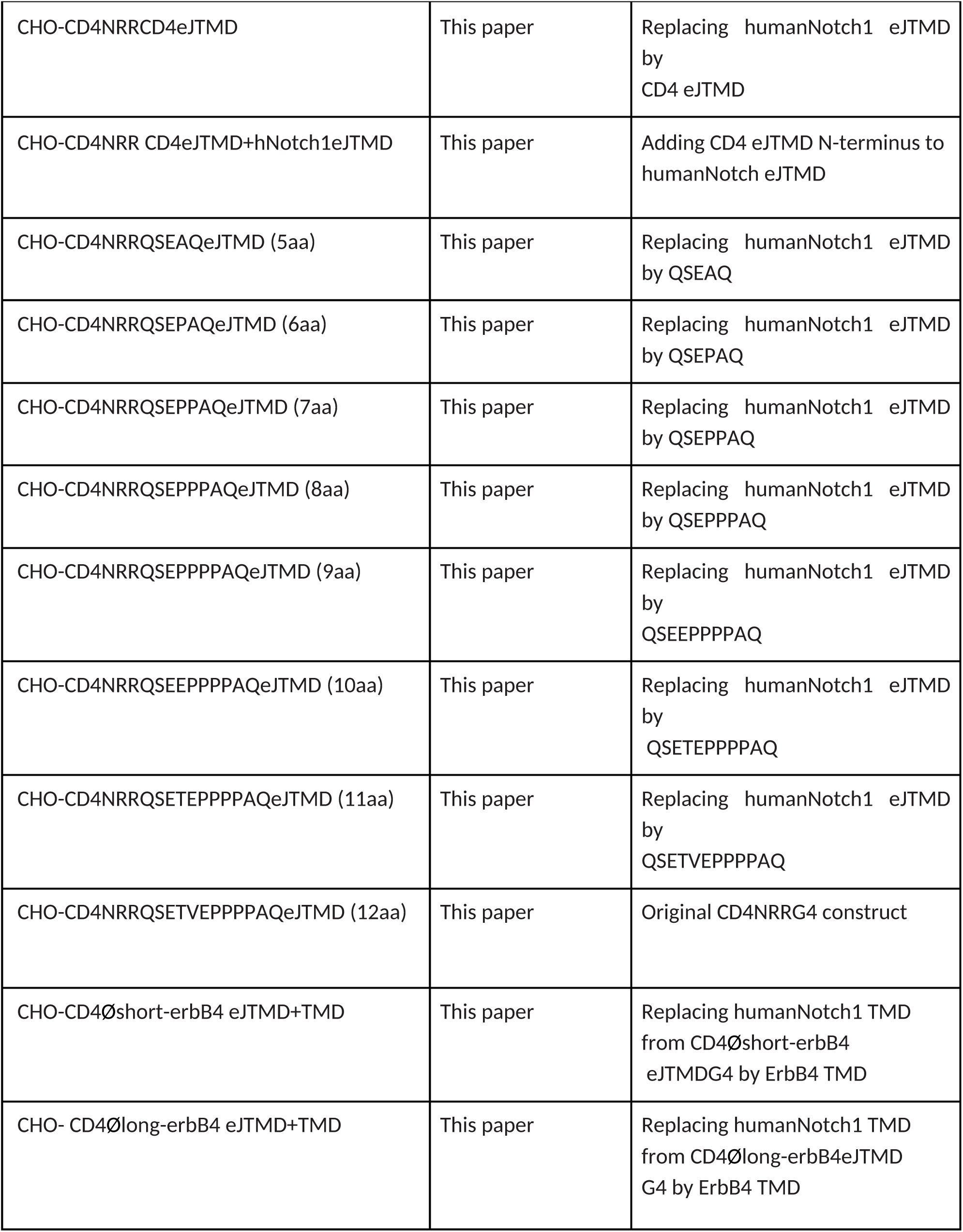
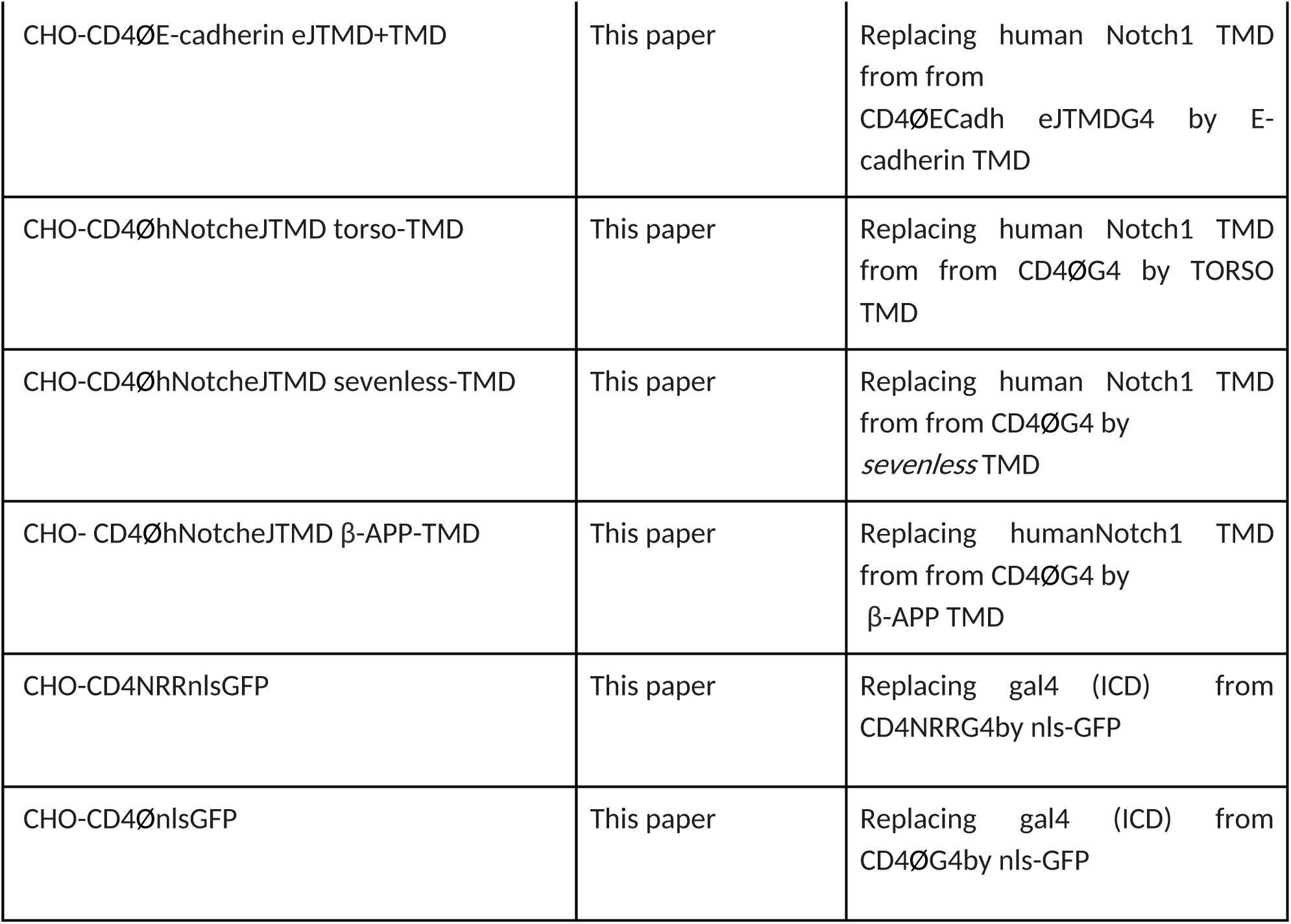

### Cell-cell interaction experiments

Receptor cell lines were co-cultured with cells expressing their cognate ligands (mCD19mCherry, gp120mCherry, hCD19tdTtomato, or hDll4) at 1:1 ratio in 24-well plates (at 300,000 cells/ml). 48 hours after plating, cells were photographed under an inverted epifluorescence microscope with a 10x objective. At least three independent replicates were made for each receptor.

### Induction experiments by substrate-attached ligands on ELISA plates

Mouse anti-CD4 (monoclonal-CD4 (MHCD0400, Thermo Fisher), mouse anti-humanNotch1 (MAB5317, R&D Systems), and anti-rabbit anti-mouse IgG F(ab’)2 (315-005-047 Jackson ImmunoResearch) antibodies were diluted at 10 μg/ml. Diluted antibodies were used to coat 96-well ELISA plates (442404, Thermo Scientific) at 4°C overnight. Next day the ELISA plates were gently washed with PBS, and 10mg/ml BSA was added to block the plates at 37°C for 2 hours. Then, receptor cell lines carrying the reporter (UAS-H2BmCitrine) were plated at (2 x 10^5^).

For S2 inhibitor experiments, we incubated the cells with a cocktail containing batimastat (BB94, 50 μM; SML0041, Sigma-Aldrich), GM6001 (50 μM; SC203979, Santa Cruz Biotechnology), and TAPI (100 μM; SC20585, Santa CruzBiotechnology). For S3 inhibitor experiments, DAPT (10 μM) was added into the growth medium when cells were plated. Cells were imaged 48 hours after plating using an inverted epifluorescence microscope with a 10x objective.

### Downregulation of ADAM10 by CRISPR-CAS9

We used CRISP/Cas9 to knockout the ADAM10 gene in human (Hela cells). We tested 3 different gRNAs against human ADAM10: (#1 346-CATGGGTCTGTTATTGA; #2 982-CTTGGTCTGGCTTGGGT; #3 1123-GTAATGTGAGAGACTTT; abm, 1130711; HGNC ID:188). We made lentiviruses, infected Hela cells and verified the efficiency of CRISP by western blot (rabbit polyclonal (abcam, ab1997, diluted 1:800). We observed that #3 gRNA was the most effective one at downregulating ADAM10 and used it for all experiments to infect the receptor cell lines CD4NRRG4 and CD4ØG4. Then, we performed induction experiments by substrate-attached ligand and images were taken after 48 hours of cells were plated.

### Imaging and quantification of induction experiments by cell-cell interaction and by substrate-attached ligand on ELISA plates

For all *in vitro* experiments, images were taken under an inverted fluorescence microscope with 10x or 20x objective lenses with an exposure time of 200 milliseconds, and 1 binning under the same conditions across constructs and experiments.

Images were acquired in uint16 format. The images were smoothed with a Gaussian filter (standard deviation 2), and divided by the same image smoothed with a Gaussian filter using a standard deviation of 3. The resulting ratiometric image was normalized by dividing each pixel by the maximum intensity in the image. The normalized image was then converted to a binary using a threshold of 0.501. The Gaussian filters and threshold size were empirically determined and kept constants for all experiments unless stated otherwise. The binary image was then segmented using the MATLAB function *regionprops* and each ROI was labelled with a unique identifier. The Euclidean distance transform of the labelled image was then calculated to obtain the interpixel distance between all pixels in an ROI. This approach assigns pixels at the core of an ROI with a higher value than pixels at the periphery. The Euclidean distance matrix was inverted, and the watershed approach was used to segment the matrix into a series of segments, some of which contain the expected ROIs. This step is necessary to capture features of the ROIs which may not be Gaussian in nature. The fully segmented image was then multiplied by the original labelled binary image and relabeled with unique identifiers. The resulting segmentation contains the non-Gaussian shape of ROIs above a threshold. The same approach was applied to the induction and background images. The ROIs from each image were pooled together, and unless stated otherwise, were analyzed as a population. The distribution of area and intensity was plotted and used to manually select areas corresponding to the size of a cell. The same area limits were used to analyze the induction and background datasets but varied across experiments (usually in the range of 10 and 2000 pixels).

The mean intensity of each ROI was calculated and summed across all ROIs in an image to obtain the integrated intensity for each image. The median, and standard deviation of the integrated intensity was calculated across all images in multiple experiments performed under identical conditions. The fold induction was obtained by calculating the ratio of integrated intensities at different conditions. The first step was to obtain the probability distribution of ROIs with a specific mean intensity. To obtain a proper sampling of all intensities, the ROIs from 5 images were pooled together. The probability distribution represents the median and standard deviation across multiple sets of 5 images each. The same approach was used for the induction, and background images. A threshold was selected by identifying a particular intensity with equal probability in both the induction and background images. In cases where the intensity probabilities did not intercept at any intensity, a threshold was selected based on the curvature of the distribution. The fold activation was calculated by dividing the integrated intensity of ROIs above the threshold in the induction images by the integrated intensity of ROIs above the same threshold in the background images. The intensity fold induction was taken across all possible induction-background image pairs in experiments performed within the same day.

In some experiments where segmentation was not possible due to cytosolic expression of the fluorescent reporter, we only determined the overall intensity of the images by summing the intensity of every pixel in the images. The fold-change in intensity in when the blockers are applied or knocking down the ADAMS gene with CRISPR-CAS9 were calculated by the following equation.

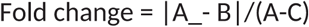

A is the summed intensity of an image with ADAMS in the ligand cell line’s presence. B is the receptor cell line with knocked down ADAMS in the presence of the ligand cell line. The letter C represents the summed image intensity of the receptor line with knocked down ADAMS in the absence of the ligand cell line. To investigate the effect of distinct manipulations on the overall intensity of induction, we divided the summed intensity of the images in the blocker’s presence by the summed intensity of the images without the blockers. Because all images were taken under identical conditions, the equation shown above was applied between randomly selected pairs of images.

### Time-lapse co-culture assays and microscopy

24 Glass-bottom multi-well plates (CLS-1812-024 from Chemglass Life Sciences) were coated with 5 μg/ml Hamster Fibronectin (Oxford Biomedical Research) diluted in 1x phosphate-buffered saline (PBS) for 30 minutes at room temperature. Emitter cells (gp160 ligand cell line) or CHO-K1 cells were mixed in suspension with similarly trypsinized receptor cell lines (CD4NRRG4 orCD4ØG4) at a ratio of 10:1. A total of 10 x 10^4^ cells (70% confluence) were plated for each experiment. Imaging starts 2-4h post-plating.

#### Time-lapse microscopy

Movies were acquired at 20X (0.75 NA) on an Olympus IX81 inverted epifluorescence microscope equipped with autofocus (ZDC2), and an environmental chamber maintaining cells at 37°C, 5% CO2. Automated acquisition software (METAMORPH, Molecular Devices) was used to acquire images every 30 minutes in multiple colours (YFP, RFP) or differential interference contrast (DIC), from multiple stage positions.

### Quantification of Time lapse of Synthetic Receptors induction *in vitro*

To investigate the time evolution of intensity changes upon cell-cell interactions, we performed live imaging of 25 cells for each construct and obtained images at 512 by 512 pixels every 25 minutes. The analysis of intensity changes across time was performed by tracking the changes in intensity upon induction of individual cells. The intensity changes of individually tracked cells was determined by manually drawing a boundary around a subset of selected cells in each image. Only cells which were visible in all frames were selected. Whenever a cell would divide into two cells one arbitrarily selected cell would be tracked for the remaining of the video. The manually drawn boundaries were refined automatically by applying the contour region growing technique (Chan and Vese, 2001) on the Gaussian filtered image (1-pixel sigma) at each frame. The refined boundary’s median intensity was calculated and averaged across tracked cells in the induction and background conditions. Individual tracking: the changes in intensities of individually tracked cells were monitored by calculating the median intensity within the refined cell boundary of a cell in each frame. The median intensity within the refined boundary as a function of time was fitted with a shifted logistic equation (eqn. 2).

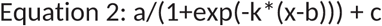

Where x is time, b is the median intensity at half-saturation, (a + c) represents the maximum median intensity, c is the initial median intensity, and K is the rate of change. The fold induction for a cell was calculated by dividing the maximum median intensity of the ROI (a+c) by the initial median ROI intensity (c). Because the median intensity of an ROI would change abruptly following a cell division event, we ignored periods after cell division. Changes in intensities during these manually selected time windows were fitted with the above equation, and the rate constant was recovered for each individually tracked cell. The median and standard deviation of the half-saturation midpoint (b) or the fold induction were pooled across trials and statistical significance was determined as described below.

### Western Blotting

#### Sample preparation

Cells were washed with PBS before adding 500 μl of RIPA (Radio Immuno Precipitation Assay) Buffer (50mM Tris, pH8.0, 150mM NaCl, 1% Triton X-100, 0.5% sodium deoxycholate, 0.1% SDS, and EDTA-free protease inhibitor cocktail) at room temperature into a 6 cm plate. Cells were scraped with the lysis buffer and incubated for 5 minutes before shearing the genomic DNA by passing the lysate 15-20 times through a 26G needle. Then, the cell lysate was either boiled for 3 minutes (ADAM10 detection) and spinned down at 4°C 14.000 rpm for 5 minutes to discard the insoluble cell debris. The lysate supernatant was transferred to a new Eppendorf tube and used immediately or stored at - 20°C until further use. If the samples were frozen, they were thawed, boiled for 2 minutes and spun down before loading.

#### Measurement of protein concentration

The protein concentration of the cell extracts is determined by the Bradford colourimetric method, using bovine serum albumin (BSA) as standard. The protein concentrations were determined by measuring the samples’ absorbance at 594 nm and extrapolating the values obtained in a standard line constructed with known BSA amounts (125-2000 μg).

Western blotting was performed using standard methods. antibodies against ADAM10 (rabbit polyclonal (abcam, ab1997, diluted 1:800), alpha-tubulin (mouse monoclonal (DM1A (ThermoFisher) diluted 1:500) overnight in 1% of blocking solution at 4°C. Next day, the membrane was washed 3 times with TBST for 5 minutes each wash and incubated with secondary antibody conjugated to the enzyme horseradish peroxidase (HRP) (goat anti-rabbit (1706515,BIO-RAD diluted 1:2000), and anti-mouse (1706516, BIO-RAD diluted 1:2000) in 1% of blocking solution for 1hour at room temperature. After the incubation with the secondary antibody, the membrane was washed 3 times for 5 minutes in the TBST buffer. Finally, the chemiluminescent substrate (1705061, BIO-RAD) was added for 2 minutes at room temperature. The membranes were imaged with a Western Blot imager (Azure c-400 from Azure system) at different exposure times.

### Live cell Immunostaining

The cells for immunostaining were first seeded on coverslip glasses pre-coated by poly-D-lysine (P6407, Sigm-Aldrich) in water for 30 minutes at room temperature. For cell surface staining, the cells were first pre-treated with Dynasore (100 μM, D7693, Sigma) for 30 minutes at 37°C in the incubator to block endocytosis and maximize the amount of surface protein. The cells were then washed by the surface staining buffer (1x PBS with 2% fetal bovine serum (FBS), 0.1% sodium azide, and 100μM Dynasore) for 5 minutes on ice. After the wash, the cells were incubated with different primary antibodies depend on the receptor cell line: anti-human CD4 (1:50 dilution, monoclonal-CD4(MHCD0400, Thermo Fisher) or rabbit anti-mouse IgG F(ab’)2 (1:50 dilution in the surface staining buffer, 315-005-047, Jackson ImmunoResearch) for 60 minutes on ice. The cells were washed three times for 5 minutes each on ice after the antibody incubation. Subsequently, the cells were incubated with the secondary antibody, goat anti-rabbit or anti-mouse (Alexa555, 1:250 dilution) for 45 minutes on ice. After this, the cells were washed three times for 10 minutes each. After the cell surface staining, the cells were fixed with 4%paraformaldehyde for 10 minutes at room temperature and followed by the regular immunostaining procedures or incubate with 499 fluor Membrite (fluorescent dye that stain and react with cell surface proteins, 30093-T, Biotium) for 30 minutes at 4°C and DAPI (to label cell nuclei 4ʹ,6-diamidino-2-phenylindole, 1:50,000) for 10 minutes at room temperature. The cells were washed in PBS three times for 5 minutes each and permeabilized with PBS/0.05% triton X-100 (PBST) for 20 minutes for immunostaining. After the permeabilization process, a blocking solution with 10% FBS in PBST was added to the cells for 40 minutes. Subsequently, the cells were incubated with different primary antibodies depending on the experiment: antibodies against Gal4DBD (DNA binding domain, mouse monoclonal IgG2a, (RK5C1) sc-510. Santa Cruz, Biotechnology diluted at 1:200) or mCherry/tdTomato (Rat monoclonal [5F8] to Red Fluorescent Proteins (RFP), Chromotek; diluted at 1:1000) diluted in 1% serum/PBST. Cells were incubated overnight at 4°C. Next day, the cells were washed three times in PBST and incubated with secondary antibody (555 goat anti-mouse or rat, Life Technologies, diluted at 1:800) for 90 minutes. Finally, the cells were washed 3 times in PBST and analyzed under an inverted epifluorescence microscope or imaged using a confocal microscope (Zeiss LSM 800) under a 63x or 100x objective.

### Biochemical identification of the cleavage site in constructs without NRR

CHO-K1 cell lines were transfected with two different receptors: CD4-hN1NRR-hN11TMD-nlsGFP or CD4-hN1Ø-hN1TMD-nlsGFP using lipofectamine 2000 reagent (Invitrogen, 11668027). The cells were seeded 24h before the transfection to be 70 - 90% confluent the day of transfection. Lipofectamine 2000 was diluted in Opti-MEM in a 1:5 ratio. The DNA was also diluted in the same medium: 24 μg of DNA in 1.5 ml of medium for 10 cm plates. Both dilutions were incubated at room temperature for 5 minutes and then mixed and incubated for at least 20 minutes at room temperature. Then, the lipid-DNA complex was added to the cells. After 4 hours, the cells were washed with fresh medium containing DAPT (γ-secretase blocker) or DMSO as a control. In order to find the S2-like site of the receptor without NRR, the S3 cleavage site must be blocked because the protein is immunoprecipitated by nanobodies that recognize the ICD. DAPT was added to the medium after 4 hours of transfection. Twenty-four hours after the washes, ligand cell lines expressing gp120mCherry or mCD19mCherry (as a negative control) were plated on top of the CHO-K1 cells transfected with different receptors. The cells were harvested after 48 hours of adding the ligand cell lines.

After co-culture induction assay, the cells were washed with DPBS (without calcium and magnesium) at room temperature and collected with a scraper. Cells in suspension were centrifuged for 5 minutes at 1000 rpm with two washes in between. The pellet was frozen at -80°C. The pellet was thawed and diluted in solubilization buffer (0.05M HEPES pH 7.5, 0.2MNaCl, 2mM MgoAc (Millipore Sigma, 63052-100), Triton 1%, 25x EDTA-free protease inhibitor cocktail (PI, Millipore Sigma, 11873580001), 1mM DTT (1,4-dithiothreitol, Millipore Sigma), and 4M UREA, and incubated at 4°C for minutes. The pellet was then centrifuged at 18,000 relative centrifugal force (rcf) for 20 minutes in a table-top Eppendorf centrifuge at 4°C.

Biotinylated nanobodies (kind gift Dr. Rebecca Voorhees [Caltech]), which target the intracellular domain of the receptors (nls GFP), were immobilized on magnetic Pierce Streptavidin Beads (Thermo Fisher) in wash buffer (0.05M HEPES pH 7.5, 0.2M NaCl, 2mM MgoAc (Millipore Sigma, 63052-100), Triton 0.1%, 25x EDTA-free protease inhibitor cocktail (PI, Millipore Sigma, 11873580001), and 1mM DTT (1,4-dithiothreitol, Millipore Sigma) for 30 minutes at 4°C. The remaining biotin-binding sites on the beads were subsequently blocked with 50 μM Biotin-PEG-COOH (Iris Biotech) for 15 minutes in solubilization buffer. The blocked beads were incubated with the Protein Extract for 1h at 4°C. After the incubation, the beads were separated from the extract using a magnetic rack and were washed twice in the solubilization buffer followed by two washes in the wash buffer. Nanobody-target protein complexes were then eluted by adding 0.5 μM SUMOStar protease (Liu et al., 2008) in wash buffer for 30 minutes at 4°C. Once the proteins were purified, digestion in solution with Chymotrypsin and trypsin was performed.

The four samples (induction (with gp120 ligand) and control (with mCd19 ligand) for CD4NRRG4 and CD4ØG4) were digested following a reduction process with 1 μl of TCEP (Tris(2-carboxyethyl) phosphine, reducing agent) in MS buffers for 20 minutes at room temperature and alkylation with 3.6 μl OF 500 mM 2-chloro-acetamide for 15 minutes at room temperature. After this, the samples were incubated with 2 μl Lys-C endoprotease for 4 hours at 37 °C. Then, the samples were incubated with trypsin (100 ng/μl), and Chymotrypsin (100ng/ μl) for 18 hours at 37°C and at 25°C for 18 hours, respectively. The next day, the samples were desalted using Pierce C18 spin columns (cat #89870). 10 μL of 20% TFA were added to the samples to adjust pH=2. Then, an activated resin process of the columns was performed. After the resin equilibration, the samples were added to the columns and centrifuge for 2 minutes at 1500 g three times. Then, the samples were washed (200 μL 5%CAN (acetonitrile), and 0.5% TFA (Trifluoroacetic acid) and eluted with 50 μL 70% ACN 0.2% FA (formic acid). Finally, the samples were dried and stored at -80°C forMass Spectrometry.

Liquid chromatography-mass spectrometry (LC-MS) analysis was carried out on an EASY-nLC 1200 (Thermo Fisher Scientific, San Jose, CA) coupled to an Orbitrap Eclipse Tribrid mass spectrometer (Thermo Fisher Scientific, San Jose, CA). Digested and desalted peptides were resuspended in 20 μL 0.2% formic acid and 5 μL peptides per sample were loaded onto an Aurora 25 cm x 75 μm ID, 1.6 μm C18 reversed-phasecolumn (IonOpticks, Parkville, Victoria, Australia), and separated over 43 minutes at a flow rate of 350 nl/min with the following gradient: 2–6% Solvent B (3 minutes), 6-25% B (20 minutes), 25-40% B (7 minutes), 40-98% B (1 minute), and 98% B (12 minutes). Solvent A consisted of 97.8% H2O, 2% ACN, and 0.2% formic acid, and solvent B consisted of 19.8% H2O, 80% ACN, and 0.2% formic acid.

MS1 spectra were acquired in the Orbitrap at 120K resolution with a scan range from 350-2000 m/z, an AGC target of 1e6, and a maximum injection time of 50 milliseconds in Profile mode. Features were filtered for monoisotopic peaks with a charge state of 2-7, and a minimum intensity of 1e4, with dynamic exclusion set to exclude features after 1 time for 45 seconds with a 5-ppm mass tolerance. HCD fragmentation was performed with collision energy of 28% after quadrupole isolation of features using an isolation window of 0.7 m/z, an AGC target of 1e4, and a maximum injection time of 35 milliseconds. MS2 scans were then acquired in the ion trap at Rapid rate in Centroid mode and with auto scan range. Cycle time was set at 3 seconds.

### Data Analysis

LCMS data was searched and quantitated using ProteomeDiscoverer 2.5 (ThermoFisher Scientific) (Lemeer et al., 2012), of which the protein database consisted of 23885 proteins from the *Cricetulus griseus* UniProt proteome (UP000001075) (“UniProt,” 2020) and 4 proteins representations of the CD4 and NRR domains. The Sequest search parameters were set to non-specific enzymatic cleavage in order to capture the non-tryptic endogenous cleavage site, peptide length between 2 and 64 residues, precursor tolerance of 20 ppm and fragment mass tolerance of 0.5 Daltons (Eng et al., 1994). Modifications were set to static carbamidomethyl on C, dynamic protein N-terminal acetylation and oxidation of M. Percolator PSM validation was set to a maximum p-value of 0.05 (Käll et al., 2007). The quantitated peptide sequence results were exported to a custom R script that assembled the peptides into protein sequences, accounted for the summed quantitation of individual amide residues and generated the visualizations provided in the discussion (“R Core Team (2020). — European Environment Agency,” n.d.).

### *In vivo* experiments with transgenic *Drosophila*

Transgenic flies with PREX-dNotchNRRgal4 and PREX-dNotchØGal4 were generated by targeted insertion into the attp P2 site (Bestgene) under the control of heat shock promoter (hsp). Embryos were collected every 30-45 minutes at 25C to maximize the chances of using embryos at similar stages of development. *Drosophila* embryos received heat shocks of 2 hours at 37C and recovered for another 2 hours at 29C. The outer chorion layer was removed to visualize both receptors’ expression pattern for live imaging. The *Drosophila* embryos were dechorionated with 50% bleach for 3 minutes under gentle agitation followed by 3 washes with dH2O. After that, the embryos were mounted in holocarbon oil (H8898 from Sigma) on a glass slide. Dechorionated embryos were imaged using a confocal microscope (Zeiss LSM 800) under an 20X objective. Z-stacks were merged and analyzed using ImageJ. All the crosses were maintained at room temperature and were repeated at least 3 times.

### Statistics

The integrated intensity and/or fold-induction were compared across constructs by calculating each construct’s median integrated intensity or fold-induction for each trial. A two-sided Wilcoxon rank-sum test was performed to test the hypothesis that each construct’s median values belonged to distinct distributions with unequal medians. Unless stated otherwise, all statistical tests were performed using nonparametric tests.

The Symbols for the p-values are indicated in the next table:

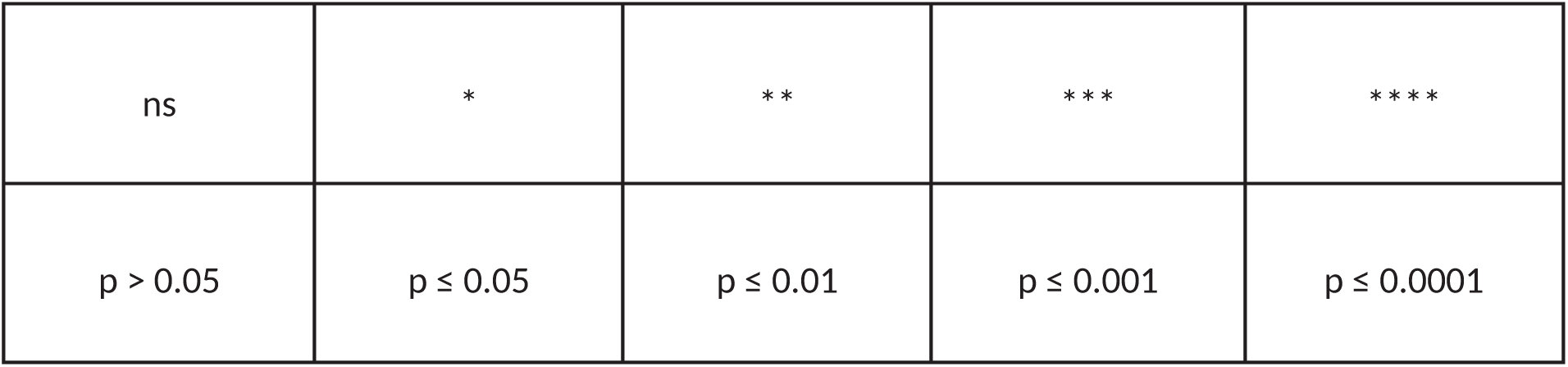

