## Supplementary material for "SHORT DISORDERED PEPTIDES ARE SUFFICIENT TO CONVERT PROTEINS INTO MECHANOSENSORS THAT RESPOND TO PHYSIOLOGICAL CELLULAR FORCES": supplemetary figures

**SUPPLEMENTARY FIGURES.**

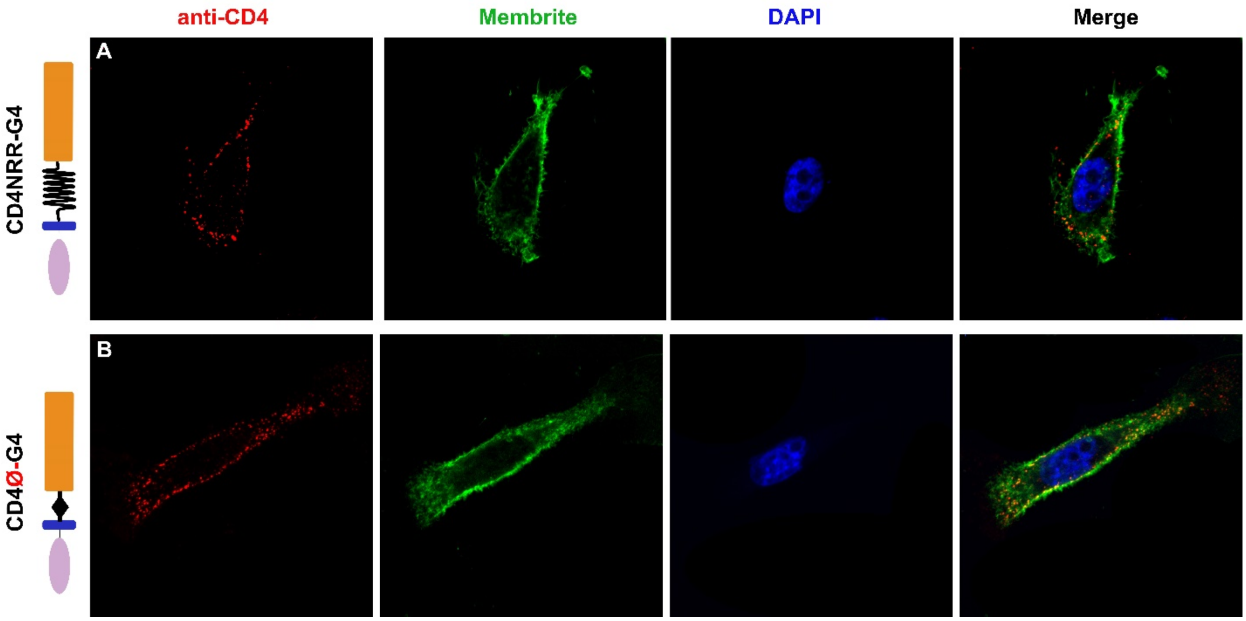


**Supplementary Figure 1. Cell membrane display of CD4NRRG4 and CD4G4. (A)** CD4 labeling the extracellular domain of CD4NRRG4 receptor did not show differences compared to CD4ØG4 **(B)**.

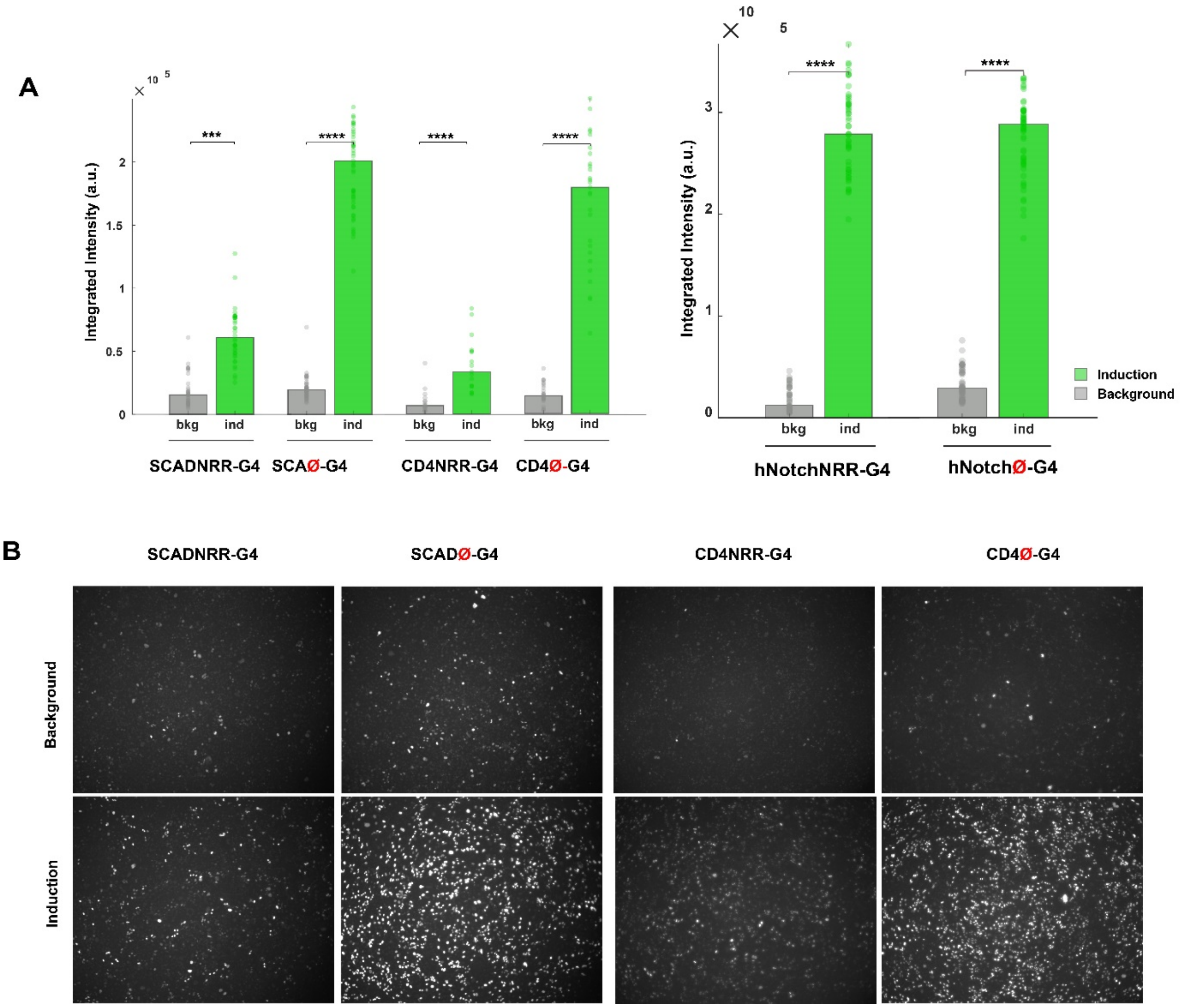


**Supplementary Figure 2**. **(A)** Integrated intensity of induction and background showing higher significant differences for engineered receptors without than with NRR, p-values between induction and background integrated intensity: hCARNRRG4=1.95 x 10-12; SCADØG4=3.12 x 10-15; CD4NRRG4=7.54 x 10-6; CD4ØG4=3.03 x 10-10; NotchNRRG4= 3.30 x $10^{-18}$ and NotchØG4=3.55 x $10^{-21}$ **(B)** Images showing the differences between induction and background for SCADNRRG4, SCADØG4, CD4NRRG4, and CD4ØG4.

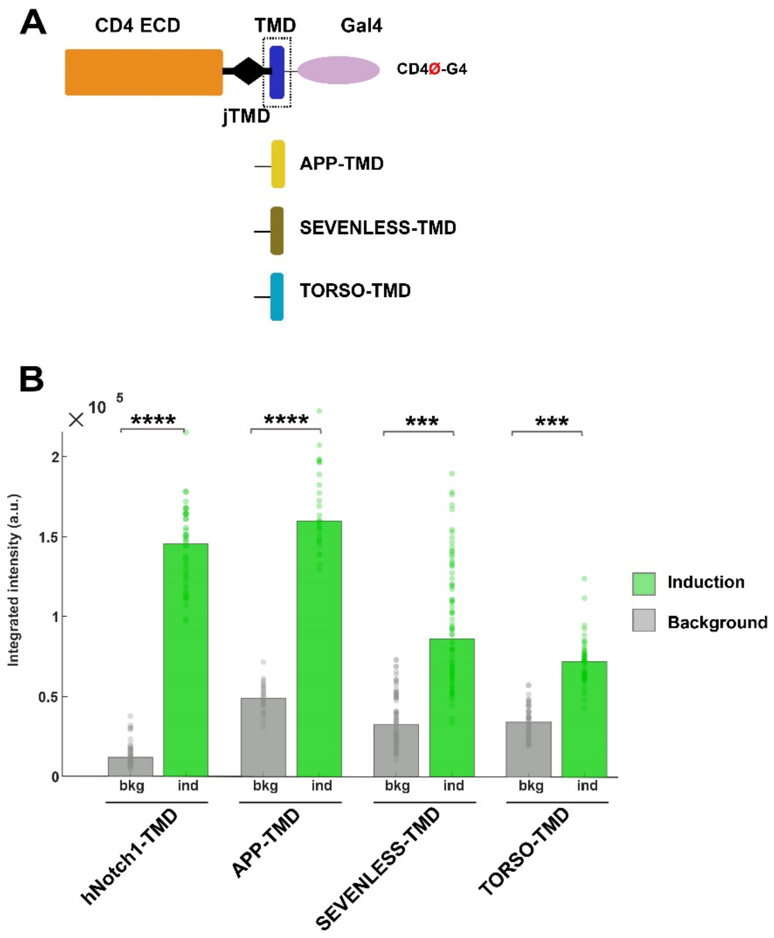


**Supplementary Figure 3. Different TMDs can be used for engineered receptors. (A)** Diagram showing TMDs from different proteins. **(B)** Integrated intensity showed induction for the three receptors carrying TMDS from torso, sevenless and APP but significantly lower than the hNotch1TMD (TORSO=2.21-fold, SEVENLESS=2.73-fold, and β-APP=3.21-fold; p-values compared to hNotch1eJTMD: TORSO=4.14 x 10-13, SEVENLESS=9.76 x 10-15; and β-APP=8.87 x 10-13).

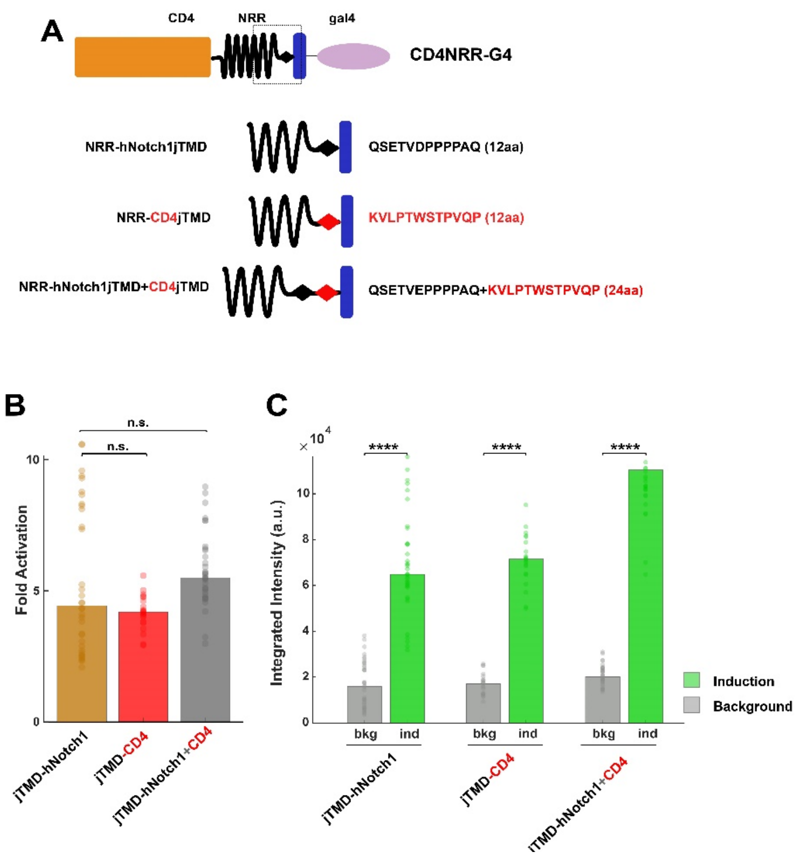


**Supplementary Figure 4. NRR lengths eJTMD. (A)** Diagram showing the amino acid sequence from CD4eJTMD and the combination of hNotch1 and CD4eJTMDs. **(B)** Fold activation did not show significant differences of both receptors compared to CD4NRRhNotch1eJTMDG4 receptor (CD4NRRhNotch1eJTMDG4=4.43-fold; CD4NRRCD4eJTMD=4.17-fold; CD4NRRhNotch1eJTMDCD4eJTMD=5.46-fold). p-values compared to CD4NRRhNotch1eJTMDG4: CD4NRRCD4eJTMD=0.520; CD4NRRCD4eJTMDhNotch1eJTMD=0.108. **(C)** Integrated Intensity graphs showed a higher level of background and induction for the receptor carrying the eJTMDs from hNotch1 and CD4.

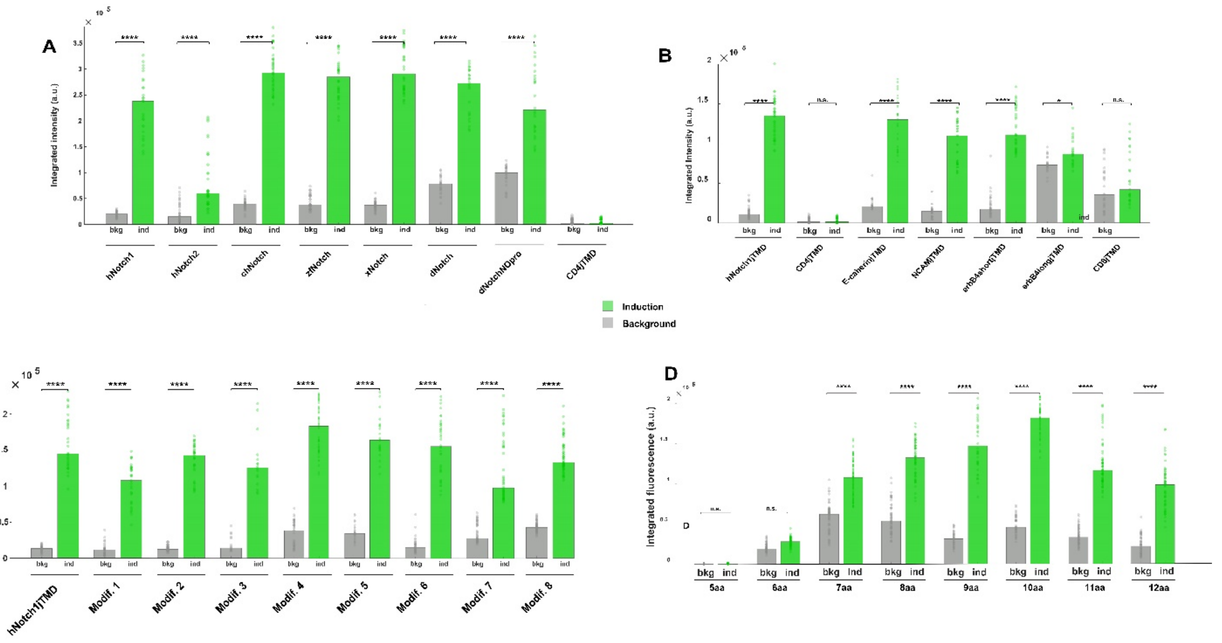


**Supplementary Figure 5.** (A) Integrated intensity of induction and background showing similar level of background for chicken, zebrafish and xenopus Notch-JTMD receptors, p-values between induction and background integrated intensity hNotch1=7.36 x $10^{-20}$, hNotch2=8.00 x $10^{-10}$, chNotch=1.06 x $10^{-13}$, zfNotch=6.60 x $10^{-14}$, xNotch=3.04 x $10^{-13}$, dNotch=3.06 x $10^{-9}$, and dNotchNOpro=6.05 x $10^{-9}$. (B) CD4eJTMD showed the lowest integrated intensity for the background and induction compared to CD8eJTMD and erbB4longeJTMD that are also not induced (p-values between induction and background, CD4eJTMD=0.2807, CD8eJTMD=0.093, and erbB4longeJTMD=0.01). Note that the induction and background for E-cadherineJTMD is similar than hNotch1eJTMD but the fold activation is lower as shown in B (p-values between induction and background hNotch1eJTMD=3.12 x 10-15, E-cadherineJTMD=3.03 x 10-3, NCAMeJTMD=7.46x10-6, erbb4shorteJTMD=4.79x10-5). (C) Similar integrated intensity levels of induction and background was observed for modif. 1(QAETVEPPPPAQ), modif. 2(QEETVEPPPPAQ), modif. 3(QSSTVEPPPPAQ), and modif. 6 (QSETVEAAAAAQ) without significant difference compared to hNotch1eJTMD. Moreover, Integrated intensity for the background and induction for modifs. 4, 5, 7, and 8 presented higher levels of background than the hNotch1eJTMD, (p-values between induction and background: hNotch1eJTMD=6.51 x $10^{-12}$; modif. 1=1.55 x $10^{-6}$; modif. 2=3.02 x $10^{-12}$; modif. 3=1.66 x $10^{-6}$; modif. 4=1.64 x $10^{-5}$; modif. 5=1.24x$10^{-5}$; modif. 6=6.60 x $10^{-14}$; modif. 7=3.23 x $10^{-17}$; modif. 8=1.62 x $10^{-20}$). (D) Integrated intensity for the different receptors with variation on the eJTMD length. Note that the background for eJTMD with 7aa and 8aa showed the highest level of integrated intensity while the lowest level was observed in the eJTMD with 5aa (p-values between induction and background: 5aa=0.41 6aa=0.004 7aa=1.43x10-8 8aa=3.06x10-9 9aa=1.51x10-9 10aa=3.47x10-9 11aa=3.06x10-9 12aa=6.63x10-9).

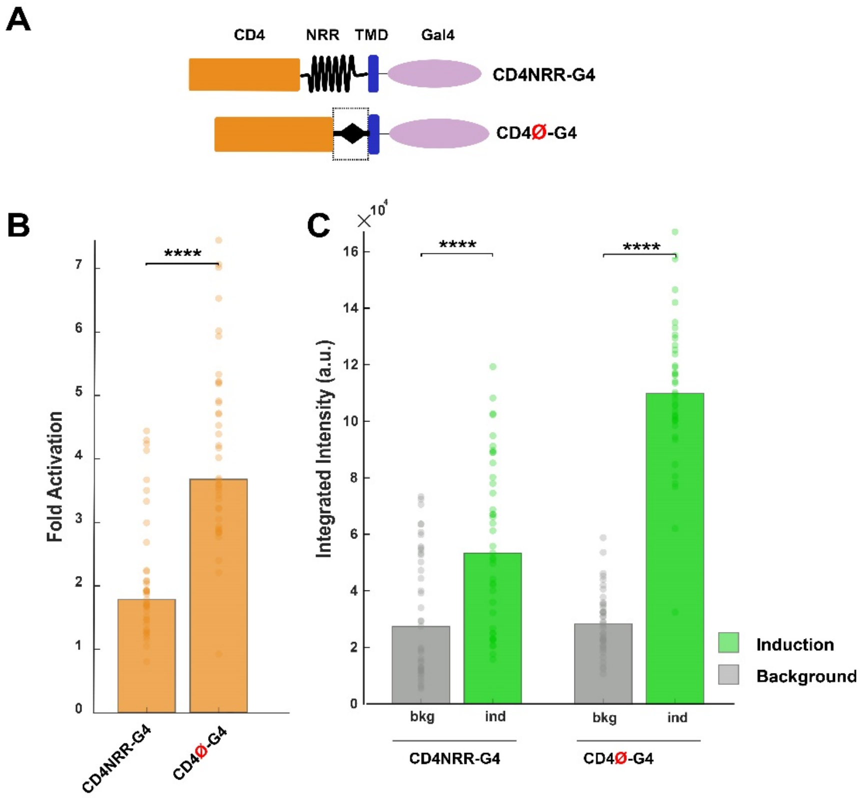


**Supplementary Figure 6. Ligand independent activation of CD4NRRG4 and CD4ØG4 receptors. (A)** Diagram showing the different domains from both receptors. **(B)** Fold increase of the CD4ØG4 (3.67-fold) receptor was higher than CD4NRRG4 (1.78-fold), p=1.37 x 10-9. **(C)** Integrated intensity of induction showed a significant difference in the level of background compared to the reporter cell line (CD4NRRG4: bkg=2.93 x 104; ind=5.33 x 104, p=5.92 x 10-5; and CD4ØG4: bkg=2.60 x 104; ind=10.98 x 104, p=4.07 x 10-14).

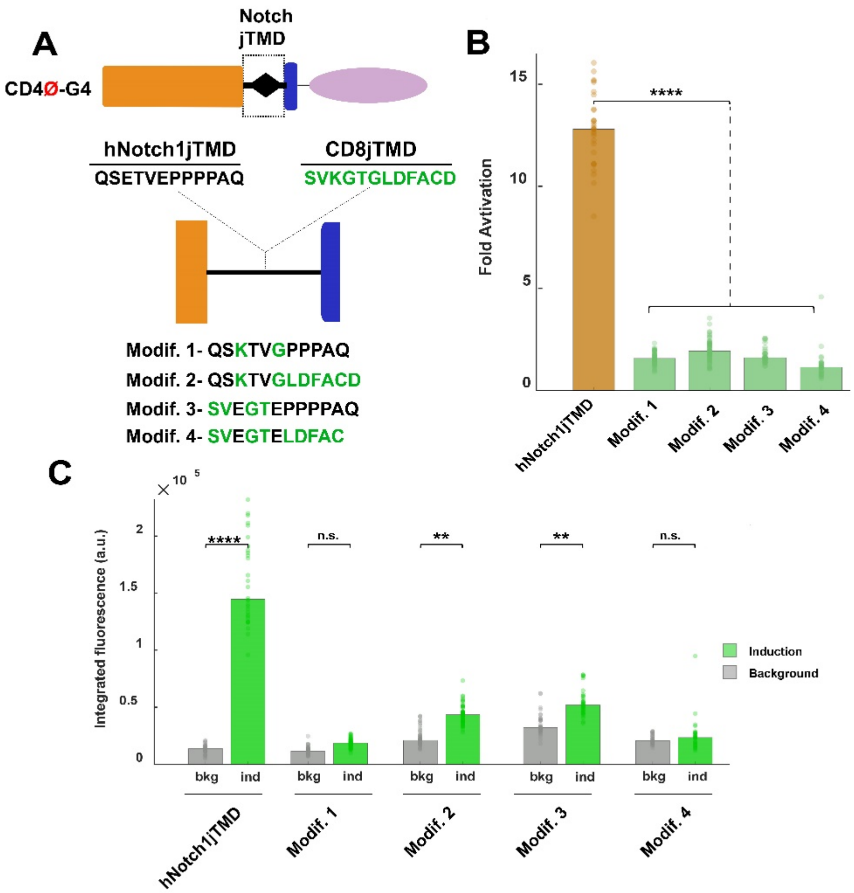


**Supplementary Figure 7. Ligand dependent induction by eJTMD modifications based on amino acids composition of CD8eJTMD and hNotch1eJTMD. (A)** Diagram showing the different eJTMD modifications based on CD8eJTMD and hNotch1eJTMD (amino acids highlighted in green are from CD8eJTMD). **(B)** Fold activation was minimal for all the modifications: modif. 1=1.57-fold, modfi. 2=1.93-fold, modif. 3=1.58-fold; modif. 4=1.12-fold. p-values compared to hNotch1eJTMD: modif1=1.50 x 10-14, modfi. 2=6.53 x 10-15, modif. 3=7.94 x 10-13, modif. 4=1.34 x 10-13. **(C)** Note that there is a small difference between the induction and background for modifications 1 and 4.
